# Haplotype Reconstruction in Connected Tetraploid F1 Populations

**DOI:** 10.1101/2020.12.18.423519

**Authors:** Chaozhi Zheng, Rodrigo R. Amadeu, Patricio R. Munoz, Jeffrey B. Endelman

**Affiliations:** Wageningen University and Research; University of Florida; University of São Paulo; University of Florida; University of Wisconsin-Madison

**Keywords:** multiparental population, hidden Markov model, tetraploid potato, double reduction, QTL mapping

## Abstract

In diploid species, many multi-parental populations have been developed to increase genetic diversity and quantitative trait loci (QTL) mapping resolution. In these populations, haplotype reconstruction has been used as a standard practice to increase QTL detection power in comparison with the marker-based association analysis. To realize similar benefits in tetraploid species (and eventually higher ploidy levels), a statistical framework for haplotype reconstruction has been developed and implemented in the software PolyOrigin for connected tetraploid F1 populations with shared parents. Haplotype reconstruction proceeds in two steps: first, parental genotypes are phased based on multi-locus linkage analysis; second, genotype probabilities for the parental alleles are inferred in the progeny. PolyOrigin can utilize genetic marker data from single nucleotide polymorphism (SNP) arrays or from sequence-based genotyping; in the latter case, bi-allelic read counts can be used (and are preferred) as input data to minimize the influence of genotype call errors at low depth. To account for errors in the input map, PolyOrigin includes functionality for filtering markers, inferring inter-marker distances, and refining local marker ordering. Simulation studies were used to investigate the effect of several variables on the accuracy of haplotype reconstruction, including the mating design, the number of parents, population size, and sequencing depth. PolyOrigin was further evaluated using an autotetraploid potato dataset with a 3×3 half-diallel mating design. In conclusion, PolyOrigin opens up exciting new possibilities for haplotype analysis in tetraploid breeding populations.

## Introduction

Polyploid species have more than two sets of chromosomes, and are especially common in flowering plants. Unveiling the genetic architecture of complex traits is fundamental in plant genetics and breeding, including for economically important tetraploid crops such as alfalfa, potato, and blueberry. Several methods have been developed for haplotype reconstruction in a polyploid bi-parental population derived from non-inbred parents (hereafter F1 population). Conditional on parental phases, Xie and Xu (2000) developed a hidden Markov model (HMM) for ancestral inference, although the model does not represent biological processes in a tetraploid F1 (Hackett 2001). Luo *et al.* (2001) developed a heuristic algorithm for parental phasing in a tetraploid F1, based on two-point linkage analyses. Hackett *et al.* (2003) modified the phasing algorithm (Luo *et al.* 2001) for analyzing SNP dosage data, and developed a HMM for ancestral inference by assuming only bivalent chromosome pairings. Zheng *et al.* (2016) developed the integrated HMM framework TetraOrigin for parental phasing and ancestral inference, accounting for both bivalent and quadrivalent formations in meiosis. The MAPpoly software (Mollinari and Garcia 2019; Mollinari *et al.* 2020) uses two-point procedures and HMMs for parental phasing and ancestral inference in polyploids up to 8 ×, assuming only bivalents.

One disadvantage of biparental populations is their limited genetic diversity, such that the discovered QTL may lose their predictive ability in a broader set of germplasm. To overcome this, many diploid multiparental populations have been recently produced, especially in crops (see review by Huang *et al.* 2015). Several software tools are available for haplotype reconstruction in diploid multiparental populations (Mott *et al.* 2000; Broman *et al.* 2003; Zheng *et al.* 2015; Broman *et al.* 2019), whereas there is no such tool for polyploid multiparental populations. The primary aim of this work is to build an HMM framework called Polyorigin for tetraploid (extendable to higher ploidy levels) multiparental populations, extending the previous framework Tetraorigin from a bi-parental F1 to multiple F1 populations that may share parents. Similar to TetraOrigin, PolyOrigin allows preferential bivalent chromosome pairing and quadrivalent formation, so that we do not make a strict distinction between allopolyploids and autopolyploids. In addition to the basic algorithm of TetraOrigin, PolyOrigin includes extra procedures to increases the robustness to the various errors in the input data.

One source of error may be the uncertainty when calling dosage from intensity signals of a SNP array or allele counts of next generation sequencing (NGS) data. We account for parental errors by a correction procedure during ancestral inference, whereas TetraOrigin introduced a parental error parameter (Zheng *et al.* 2016). In addition, we include a procedure for marker deletion during parental phasing, and the markers with parental errors are likely to be removed. On the other hand, since it has been shown that read depths of 60-80 are required for accurately inferring dosage in autotetraploids (Uitdewilligen *et al.* 2013; Matias *et al.* 2019), PolyOrigin can account for the dosage uncertainties by using NGS read count data directly.

Another source of errors is the input marker map. The marker deletion procedure during parental phasing can also remove those markers that are misgrouped or long-range misordered, in addition to parental errors. The map construction packages such as MAPpoly (Mollinari and Garcia 2019; Mollinari *et al.* 2020) and polyMapR (Bourke *et al.* 2018) order markers by the multidimensional scaling algorithm (Preedy and Hackett 2016), based on two-point linkage analyses. Such input genetic maps can be improved by an extra step of map refinement using a multi-locus HMM approach. The map refinement consists of local marker reordering and inferring inter-marker genetic distance—the latter becoming necessary when the input marker map is a physical map.

We evaluate PolyOrigin by extensive simulation studies and with a real tetraploid potato dataset. For the simulation studies, we compare PolyOrigin with TetraOrigin and MAPpoly and investigate the effect of mating design such as the number of parents. We also investigate the robustness to low depth sequencing and errors in the input dosage data and marker map.

## Methods

Figure 1 shows an overview of PolyOrigin. Suppose that we have a collection of tetraploid F1 populations. Each F1 population can be either a cross between two parents or a self-fertilization population from a single parent. The set of populations can be represented by an un-directed graph (e.g. Figure 1A), with nodes representing parents and edges representing the crosses or selfings. This is called a connected F1 since two populations can be connected by parent sharing. PolyOrigin requires two inputs: (1) a mating design describing the parents of each F1 offspring, and (2) a genotypic data matrix for all parents and offspring at a set of SNP markers. Genotypic data include a genetic map or physical map of the markers. We assumed that all markers are bi-allelic, and denote the two alleles by 1 and 2 and define a genetic dosage as the count of allele 2. We model marker data independently across linkage groups, and thus describe the model for only one linkage group.

**Figure 1:**
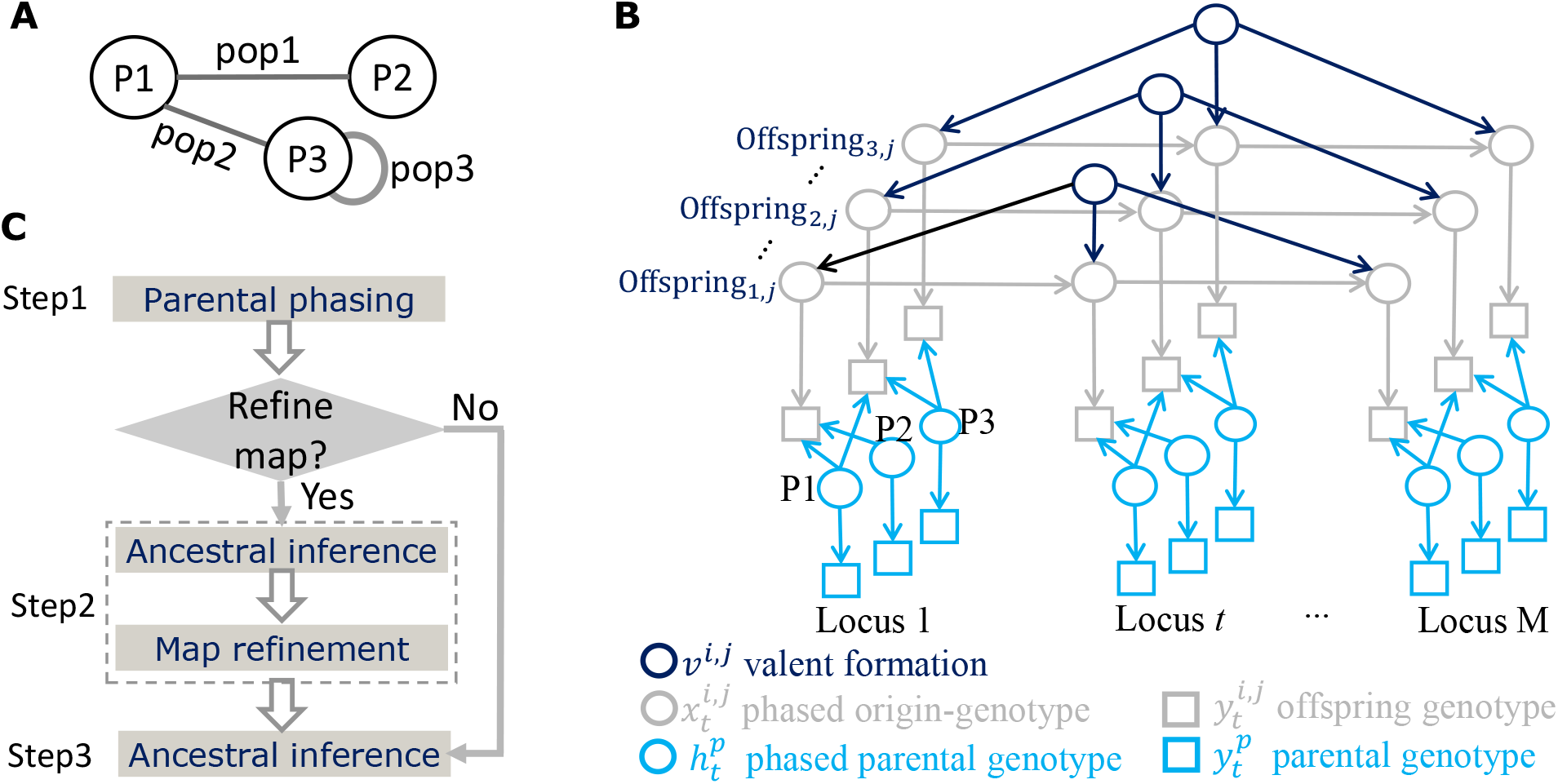
Model and workflow of PolyOrigin. (**A**) Mating design of the three F1 populations derived from three parents: P1, P2, and P3, where population 3 was derived by self-pollinating P3. (**B**) The directed acyclic graph of the PolyOrigin model for the connected F1 populations in (A). The symbol *Offspring_i, j_* denotes an offspring *j* of population *i*. The squares denote the input marker data, the circles denote random variables to be inferred, and the arrows denote probabilistic relationships to be modeled. This panel is adapted from Figure 1 of Zheng *et al.* (2016). (**C**) Workflow consists of three steps. The purpose of ancestral inference in the optional Step2 is to correct parental errors and exclude outlier offspring.

Notations for the PolyOrigin model will be introduced in the following description and are summarized in Table 1. We use *t* to index a marker, *p* for a parent, *i* for an F1 population, and *j* for an individual in a given F1 population. Denote by 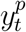 the observed genotypic data for parent *p* = 1,…, *L* at marker *t* = 1,…, *M*, and 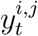 the genotypic data for individual *j* of F1 population i at marker *t*. Figure 1B shows that the PolyOrigin model has three kinds of hidden variables: 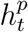 denotes phased genotype for parent *p* at locus *t*, 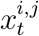 denotes phased origin-genotype for offspring (*i, j*) at marker *t*, and 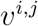 denotes valent formation for producing offspring (*i,j*) from their parents. Here the term origin-genoype denotes a combination of parental origins, referring to each parental homolog as a distinct allele.

**Table 1:** List of symbols used in PolyOrigin and their brief descriptions.

| Symbol | Description |
| --- | --- |
| $t$ | Subscript for a marker |
| $p$ | Superscript for a parent |
| $(i, j)$ | Superscript for an offspring, individual $j$ of F1 population $i$. |
| $L$ | Number of parents |
| $N$ | Total number of offsprings in connected F1 |
| $M$ | Number of markers |
| $D$ | Average number of sequence reads for an individual at a marker. |
| $S$ | Number of selfing populations in a mating design |
| $K$ | Poidy level, $K = 4$ for tetraploid |
| $P(x), P(y\vert x)$ | Probability of $x$, conditional probability of $y$ given $x$ |
| $y_t^p, y^p$ | Observed genotypic data of parent $p$ at marker $t$, $y^p = \{y_t^p\}_{t=1}^M$ |
| $y_t^{i,j}, y^{i,j}$ | Observed genotypic data of offspring $(i, j)$ at marker $t$, $y^{i,j} = \{y_t^{i,j}\}_{t=1}^M$ |
| $h_t^p, h_p$ | Hidden phased genotype of parent $p$ at marker $t$, $h^p = \{h_t^p\}_{t=1}^M$ |
| $x_t^{i,j}, x^{i,j}$ | Hidden origin-genotype of offspring $(i, j)$ at marker $t$, $x^{i,j} = \{x_t^{i,j}\}_{t=1}^M$ |
| $v^{i,j}$ | Hidden valent formation of offspring $(i, j)$ |
| $\varepsilon_t, \varepsilon$ | Genotyping error probability at marker $t$, $\varepsilon = \{\varepsilon_t\}_{t=1}^M$ |
| $\epsilon$ | Sequencing read error probability |
| $h^{\Omega(i,j)}$ | $\{h^p\}_{p \in \Omega(i,j)}$ for the parents $\Omega(i, j)$ of offspring $(i, j)$ |
| $d_t^{i,j}$ | True dosage of offspring $(i, j)$ at marker $t$, $d_t^{i,j} = f(x_t^{i,j}, h^{\Omega(i,j)})$ |
| $d$ | Genetic distance in unit of Morgan |
| $r_{bi}$ | Recombination fraction for bivalent pairing, $r_{bi} = \frac{1}{2}(1 - e^{-2d})$ |
| $r_{quad}$ | Recombination fraction for quadrivalent formation, $r_{quad} = \frac{3}{4}(1 - e^{-4d/3})$ |
| $l_t^{i,j}$ | Individual likelihood at marker $t$, $l_t^{i,j} = P(y_t^{i,j} \vert x_t^{i,j}, \varepsilon_t)$ |
| $logl$ | Marginal log-likelihood, $logl = \sum_{i,j} \log [P(y^{i,j} \vert h^{\Omega(i,j)}, v^{i,j}, \varepsilon)]$ |
| $T$ | Temperature in the simulation annealing for refining local ordering |
| $P(x_t^{i,j} \vert y^{i,j}, \varepsilon)$ | Posterior probability of phased origin-genotype $x_t^{i,j}$ |
| $P(z_t^{i,j} \vert y^{i,j}, \varepsilon)$ | Posterior probability of unphased origin-genotype $z_t^{i,j}$ |
| $f_{exch}$ | Fraction of long-range disturbed markers |
| $\sigma_{local}$ | Intensity of local disturbances in marker ordering |

The workflow of PolyOrigin consists of three steps: parental phasing, map refinement, and ancestral inference (Figure 1C). In the third step, HMM decoding and parental error correction are iterated until no errors can be detected, which are also performed prior to map refinement. The parental phasing corresponds to the maximum likelihood estimation of 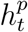. And the HMM decoding corresponds to the estimation of 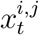, averaging over all possible 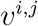 values. In the following, we will describe the basic HMM and the three steps.

## HMM

Conditional on phased parental genotypes, offspring are independent of each other. For a given offspring (*i,j*) and its valent formation *v^i,j^*, genotypic data 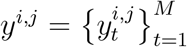 can be modeled by a HMM, which can be described by a genotype model specifying the probability of 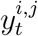 given hidden 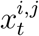, conditionally independent among markers, and a parental origin process specifying the joint prior probability of 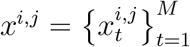

### Genotype model

At a locus t, the genotype likelihood 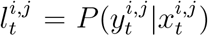 depends implicitly on parental phases via the unknown true dosage 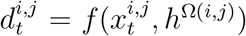, a deterministic function of hidden origin-genotype 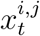 and phased genotypes 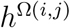 for the parents Ω(*i,j*) of offspring (*i,j*). We consider three possible representations of genotypic data 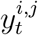. First, 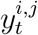 is represented by a dosage. The dosage likelihood is given by

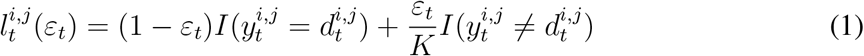

where ploidy level *K* = 4, indicator function *I*(*s*) equals 1 if statement *s* is true and 0 otherwise, and *ε_t_* denotes the dose error probability at marker *t*. If a dosage error occurs, the resulting dosage is randomly drawn from the other *K* dosages.

Second, 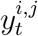 is represented by a pair of read counts. Let *r*_1_ and *r*_2_ be the counts of sequence reads for alleles 1 and 2, respectively, at marker *t* for offspring (*i,j*). Assume that the read counts *r*_1_ and *r*_2_ are generated by an unknown dosage *d′* that is different from 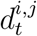 with error probability *ε_t_*, for example, because of the misalignment of reads to the reference genome. We integrate out d’ to obtain the read count likelihood

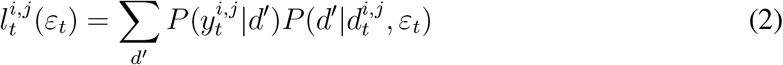

where 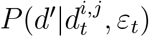 can be obtained from equation (1) by replacing 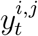 with *d′*, and 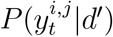 can be obtained from the following binomial model

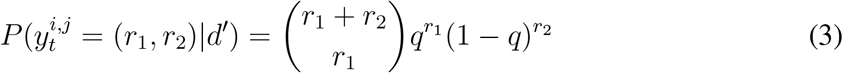

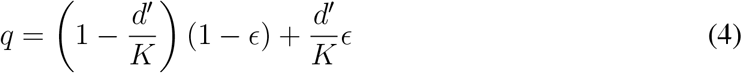

where *q* denotes the probability of a sampled read being allele 1, *d’/K* denotes the probability of allele 2 being sampled, and e denotes the sequencing error probability of observing the incorrect allele. By default, we set *ϵ* = 0.001, and the dependence of likelihood on *ϵ* is not shown in equation (3).

Third, 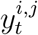 is represented by the vector of probabilities 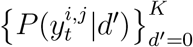, a generalization of the first and second representations, and the data likelihood is given by

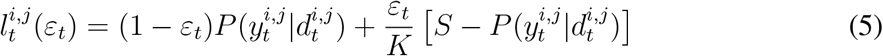

according to equations (1) and (2), where 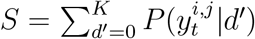. The probability vector can be calculated from equations (3) and (4) for the NGS read counts.

For example, suppose that 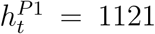 and 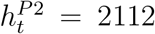 for the two parents Ω(*i, j*) = (*P*1, *P*2) of offspring (*i,j*), and 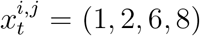 denotes that the offspring is descended from homologs 1 and 2 of parent *P*1 and homologs 6 and 8 of parent *P*2; we denote the four homologous chromosomes of the first parent *P*1 by 1 – 4, and 5 – 8 for the second parent *P*2. Thus the true phased genotype is 1112 and the true dosage is 1. If dosage 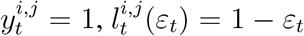. If read count 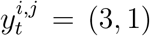, the probability vector is (0.0040, 0.4219, 0.25, 0.0471,0.0000) for *ϵ* = 0.001, and thus 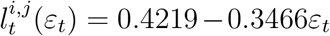. If probability vector 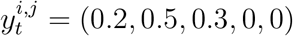, 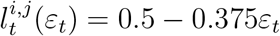. If 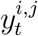 is a missing value, 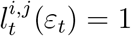.

### Parental origin process

Zheng *et al.* (2016) have described a discrete time Markov chain model for the parental origin process along four homologs of an offspring in a F1 population. The same model can be used for an offspring resulting from selfing, except that the state space is different. A discrete time Markov chain model consists of two components: a discrete distribution *P*(*x*_1_) of the states at the first marker *t* =1, and a transition probability matrix *P*(*x*_*t*+1_ |*x_t_*) describing how the states change from marker t to the next *t* +1 for *t* = 1,…, *M* – 1, so that the joint prior distribution is given by 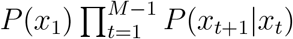, because of the Markov approximation. Here we summarize the two components.

The two gametes in an offspring are assumed to be produced independently. The initial distribution for a zygote can be obtained by the Kronecker product between the two initial distributions, one for each of the two gametes. Similarly, the transition probability matrix for a zygote can be obtained by the Kronecker product between the transition probability matrices for the two gametes. Denote by *v^i,j^* = (*v*_1_, *v*_2_) the valent formation *v*_1_ (*v*_2_) for the first (second) gamete in an offspring (*i,j*). We describe the parental origin process in a gamete, for example, the first gamete, conditional on a given value of *v*_1_.

Denoting the four homologs of the gamete parent by 1 – 4, *v*_1_ can take four possible values: [1, 2][3, 4], [1,3][2, 4], [1, 4][2, 3], and [1,2, 3, 4], where the first three values denote bivalent formations, and the last value denotes quadrivalent formation. For example *v_1_* = [1, 2][3,4], the initial distribution is assumed to be discrete uniform among gamete states (1, 3), (1,4), (2, 3), and (2,4), The transition probability matrix is given by the Kronecker product *P_bi_* ⊗ *P_bi_*, where

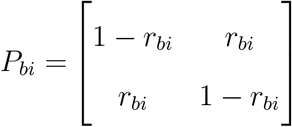

describes the transition between origins 1 and 2 along the homolog produced by the parental homolog pair [1, 2], and it refers to the transition between origins 3 and 4 for the homolog pair [3, 4]. Here *r_bi_* denotes the inter-marker recombination fraction assuming bivalent formation. For quadrivalent formation *v*_1_ = [1, 2, 3,4], the initial distribution is assumed to be discrete uniform among the 16 possible pairs of origins 1-4, and the transition probability matrix is given by *P_quad_* ⊗ *P_quad_,* where

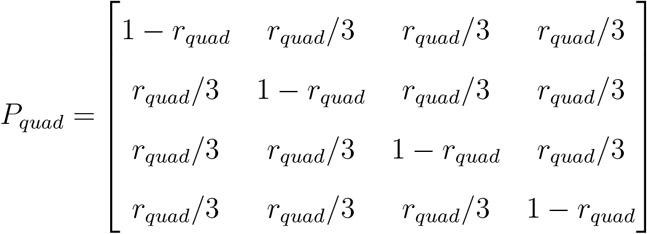

describes the transition among origins 1-4 along each homolog produced by quadrivalent formation. Here *r_quad_* denotes the inter-marker recombination fraction assuming quadrivalent formation. We assume that there is no genetic interference, and use the Haldane’s map function (Haldane 1919; Luo *et al.* 2006),
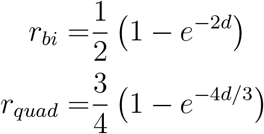

where *d* is the inter-marker genetic distance in Morgan.

If an offspring is produced by crossing between two different parents, the bivalent pairing *v*_2_ takes possible values: [5,6][7,8], [5, 7][6,8], [5,8][6, 7], and [5,6, 7,8]. If the offspring is self-fertilized, *v*_2_ takes the same set of values as those of *v*_1_. The HMM state space for a selfing offspring is thus different from that of a cross-fertilized offspring, but the size of the state space and the transition probability matrix are the same.

### Parental phasing

We extend the phasing algorithm of Zheng *et al.* (2016) from a single bi-parental F1 cross to connected F1 populations. The phasing algorithm is to optimize the log-likelihood *logl* = ∑_i,j_ *logl^i,j^*, where the individual log-likelihood 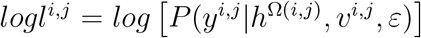. Here 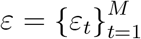 for genotyping error probabilities at all markers, 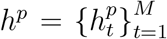 for the hidden phased genotypes of parent *p* at all markers, while the hidden origin-genotypes 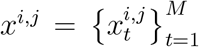 are integrated out in *logl^i,j^*. Note that *logl* depends implicitly on the marker ordering and intermarker distances.

The phasing algorithm starts with the initialization of 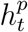 for all parents by randomly drawing 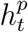 from its prior distribution 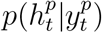. For example, if dosage 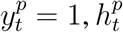 follows a prior uniform discrete distribution among the four possible phased genotypes: 1112, 1121, 1211, and 2111. If probability vector 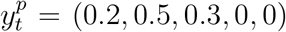, 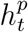 takes 1111 with probability 0.2, takes one of the four phased genotypes: 1112, 1121, 1211, and 2111 with equal probability 0.125, and takes one of the six phased genotypes 1122, 1212, 1221, 2112, 2121, and 2211 with equal probability 0.05. If 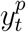 is a pair of read counts, it can be firstly transformed into a probability vector. If 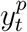 is a missing value, 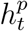 takes one of the 2^*K*^ = 16 phased genotypes with equal probability 1/16.

After initialization, each phasing iteration performs alternative maximization among valent formations and phased parental genotypes. First, independently for each offspring, the valent formation *v^i,j^* is given by maximizing the individual log-likelihood *logl^i,j^* with respect to *v^i,j^*, conditional on the phased parental genotypes *h*^Ω(*i,j*)^. For the sake of computational efficiency, we consider only bivalent formation. We calculate the individual log-likelihood *logl^i,j^* for a given *v^i,j^* by the forward algorithm for HMM (Rabiner 1989). Second, sequentially for each parent *p* = 1, …,*n_p_*, we obtain the maximum possible *h^p^*, conditional on valent formations {*v^i,j^*}_*i,j*_· for all offspring and phased genotypes (*h*^*p′*^}_*p′≠p*_ for all the other parents. Specifically, we calculate a proposed phase *h^p^* that approximates the maximum possible phase, accept it if the target function *logl* is increased, and otherwise reject it and keep the current phase. We obtain proposed *h^p^* in a forward-backward procedure, which can be adapted from the detailed description for a single F1 population (Zheng *et al.* 2016).

When phasing iteration gets stuck such that the proposed parental phase for every parent is rejected, we delete markers that do not fit into the marker sequence. Because the number of markers deleted is negatively correlated with genotyping error probability, we estimate *ε* by maximizing the target function *logl*, prior to marker deletion, assuming that *ε* does not vary with markers. We perform the estimation of *ε* and marker deletion only once for the sake of computational efficiency. We delete markers using the Vuong’s closeness test, a likelihoodratio-based test that can be used for comparing two non-nested models (Vuong 1989). We calculate the Vuong test statistic for all markers simultaneously and delete those markers with p-values significant at 0.05.

A single phasing run stops if the parental phases do not change for 5 consecutive iterations, or the number of iterations reaches 30. To find the global maximum, we perform multiple phasing runs independently and select the one with the largest *logl*. We repeat phasing runs until the so-far maximum phases have been obtained 3 times or the number of runs reaches 10. In comparison with the TetraOrigin algorithm (Zheng *et al.* 2016), we decrease some default values such as the maximum number of phasing runs, because the differences among phasing runs may be caused by the parental errors, and the PolyOrigin algorithm has additional error correction in the ancestral inference.

### Map refinement

Prior to map refinement, ancestral inference with parental error correction is performed to correct parental phase errors and exclude outlier offspring. conditional on the phased parental genotypes, map refinement iteratively updates local marker ordering, inter-marker genetic distance, valent formation *v^i,j^*, and marker-specific error probability *ε_t_*. The estimation of *v^i,j^* is the same as that in the parental phasing, except that quadrivalent formation is allowed. To decrease the effect of offspring genotyping errors, *ε_t_* is estimated by maximizing logl using the local Brent method (Brent 1973), sequentially for marker *t* =1, …,*M*, and markers with *ε_t_* ≥ 0.5 are deleted. Similarly, inter-marker distance is estimated by maximizing *logl* using the local Brent method (Brent 1973).

In each iteration, the local marker ordering is refined by sliding a window along chromosome at a step of one marker, and the ordering refinement starts with window size 2 and increases until no proposed reversion at the given window size is accepted during a scan along chromosome. The ordering of markers within a sliding window is reversed with probability min(1, *e*^Δ*logl/T*^), where Δ*logl* is the increase of *logl* due to reversion and *T* is temperature in the simulated annealing (Kirkpatrick *et al.* 1983). The temperature *T* is set to 4 in the first iteration, and decreases by half after each iteration.

The map refinement can be divided into three stages with decreasing number of updating variables. The first stage updates local ordering, inter-marker distance, *v^i,j^*, and *ε_t_*, and it changes into the second stage when *T* ≤ 0.5 and the maximum sliding window size equals 2. The second stage consists of two iterations: it updates only inter-marker distance and strips markers at a chromosome end if there exists a distance jump greater than 20 cM and the fraction of markers deleted is less than 5%. The third stage estimates inter-marker distances for selected skeleton markers in five iterations. The chromosome is divided into 50 segments, and the marker with smallest ε*t* is selected in each segment. The inter-marker distances in the final map are re-scaled piece-wisely, based on the estimated skeleton marker map.

### Ancestral inference

conditional on phased parental genotypes and the refined genetic map, each offspring is analyzed independently with a HMM. The step of ancestral inference performs iteratively the estimation of marker specific *ε_t_*, HMM decoding, and parental error correction, until there are no error corrections. The estimation of 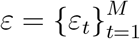 is conditional on valent formations {*v^i,j^*}_*i,j*_ for all offspring, and the estimations of *ε* and *v^i,j^* are the same as those in map refinement.

In the HMM decoding, the posterior probability 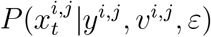 and the individual marginal likelihood *P*(*y^i,j^*|*v^i,j^,ε*) are calculated by the forward-backward algorithm for HMM (Rabiner 1989), conditional on each of the 16 possible values of *v^i,j^*, allowing for quadrivalent formation. Assuming a discrete uniform prior distribution of *v^i,j^*, we can obtain the posterior distribution *P*(*y^i,j^*|*v^i,j^,ε*) from the individual marginal likelihood according to the Bayesian theorem (Gelman *et al.* 2013). Finally, we can obtain phased origin-genotype probability

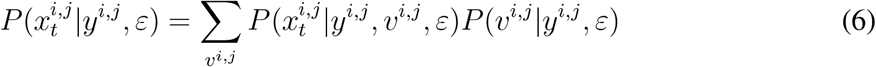

where the summation is over the 16 possible values of *v^i,j^*, and the dependencies on phased genotypes *h*^Ω(*i, j*)^ for the parents of offspring (*i,j*) are not shown.

In the parental error correction, we first perform dosage calling based on the HMM decoding. Specifically, we calculate the dosage posterior probability 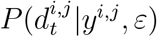 by summing the condition probability 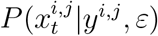 inequation (6) over 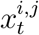 such that 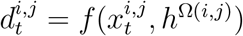. The dosage is called to be the maximum possible one if its posterior probability is larger than 0.5, and otherwise it is set to missing. Secondly, we detect suspicious markers at which the fraction of mismatches between called genotypes and observed offspring genotypes is larger than 0.15. Here mismatch refers to the input dosage being different from the called dosage, or the input probability of the called dosage being less than 0.01. Lastly, at each suspicious marker *t* and for each parent *p*, we replace the current value of 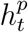 by the one with minimum mismatches in offspring dosages, among all the 16 possible values of 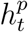, if the number of mismatches is decreased by at least 3.

The final output of ancestral inference is given by unphased origin-genotype probability 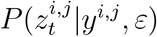 for all offspring at all markers by summarizing the corresponding phased origingenotype probabilities 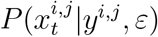, where 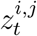 is given by the sorted value of 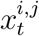. For example, unphased origin-genotype 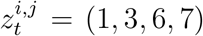 for cross-fertilized offspring (*i,j*) corresponds to four phased origin-genotypes 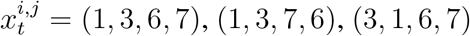, and (3,1,7, 6).

In addition, we detect outlier offspring according to the estimated distribution of the number of recombination breakpoints. Specifically, for each offspring at each marker, the unphased origin-genotype is called to be the maximum possible one if its posterior probability is larger than 0.6, and otherwise it is set to missing. For an offspring, we count the number of changes in origin-genotype along the four homologs of a linkage group after skipping the missing genotype calls, and obtain the number of recombination breakpoints by summing the number of changes over all linkage groups. An offspring is labeled as outlier if *A* > *Q*_3_ + fence * (*Q*_3_ – *Q*_1_), where the Anscombe transform 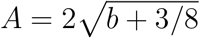 with *b* being the number of breakpoints in the offspring (Anscombe 1948), the Tukey’s fence is set to 3 (Tukey 1977), and *Q*_1_ and *Q*_3_ are the lower and upper quartiles of the transformed values.

### Algorithm evaluation

We evaluated the performance and robustness of PolyOrigin by extensive simulations using PedigreeSim (Voorrips and Maliepaard 2012) and updog (Gerard *et al.* 2018) with a custom-made R package wrap-up called PedigreeSimR available at https://github.com/rramadeu/PedigreeSimR. We quantified parental phasing error as the fraction of estimated parental phases different from the true phases, and ancestral inference error was defined as 1 minus the posterior probability of the true unphased origin-genotype, averaged over offspring and markers.

We first set up default parameter values as a baseline and then simulated four scenarios, where a few parameters varied while keeping the others at the baseline. For a given set of parameter values, we simulated three replicates and obtained results by averaging over them. **Baseline setup:** We simulated only one linkage group and first specified the true parental haplotypes. In the scenarios with fixed number of markers, the true parental haplotypes were given by the 32 real potato haplotypes; see the description in *Real Potato datasets.* The genetic length is 149 cM, with the number of polymorphic markers varying from *M* = 201 in the first two parents (*L* = 2) to *M* = 258 for *L* = 8. In the scenarios with varying number of markers, the true parental haplotypes were obtained by first simulating a genetic map and then phased parental genotypes at each marker. The inter-marker distances were first simulated from a Poisson distribution and then re-scaled to obtain the total genetic length of 100 cM, and the four homologous haplotypes of a parent were simulated by first randomly sampling a dosage and then randomly sampling a phased genotype compatible with the sampled dosage, independently at each marker.

We simulated two kinds of polysomic inheritance: (1) both preferential bivalent pairing and quadrivalent formation were allowed, *pref Pairing* = 0.5 and *quadrivalents* = 0.5, so that double reduction is possible; (2) only random bivalent pairing was allowed, *pref Pairing* = 0 and *quadrivalents* = 0, so that double reduction is not possible.

The true offspring genotypes were obtained by combining true founder haplotypes and simulated inheritance, from which observed genotypic data were obtained by applying an error model and a missing pattern. For SNP array dosage data, an error occurred in each parental or offspring dosage with probability *ε* = 0.01, and the resulting dosage was set to one of the other dosages with equal probability. Each parental or offspring dosage was missing with probability 0.1. NGS data were simulated with average depth *D* = 5, 10, …, 80, sequencing error rate 0.005, allelic bias 0.7, and over-dispersion 0.005 (Gerard *et al.* 2018). A read depth equaled zero (i.e. missing data) with probability 0.1 and otherwise followed a Poisson distribution with mean *D*/0.9.

The default mating design was a half-diallel design with *L* = 5 parents, where all 10 possible combinations of parents were crossed, and each cross produced an equal number of offspring.

#### Simulation scenarios

We divided simulated scenarios into four groups according to their study purposes: (1) comparisons with previous methods, (2) effect of population design, (3) effect of genotyping design, and (4) robustness to errors in the marker map.

To compare with MAPpoly (Mollinari and Garcia 2019; Mollinari *et al.* 2020) and TetraOrigin (Zheng *et al.* 2016), we simulated bi-parental F1 populations. Missing dosages in parents were not allowed, which is required by MAPpoly. We simulated SNP array data with population size varying from *N* =10 to *N* = 200 and two kinds of polysomic inheritance: one with double reduction and the other without double reduction.

To study the effect of population design, we simulated SNP array data for four mating designs: linear design where each parent was crossed with the next, circular design differing from the linear design by an extra cross between the first and the last parents, star design where the first parent is crossed with each of the other parents, and diallel design where all pairs of parents were crossed. The naming of mating design is based on the un-directed graph representation of the connected F1 populations. We varied three parameters: the number *L* of parents, the number *S* of selfing populations, and the total population size *N*, one at a time, while keeping all other parameter values at the baseline. When increasing *S* from 1 to 5, the selfing population was created in order from parents 1 to 5.

To study the effect of genotyping design, we simulated genotyping by SNP array and NGS data in the diallel designs with no selfings (*S* = 0), using simulated true parental haplotypes with various marker densities. The SNP array design aimed to study the robustness to genotyping error probability *ε* for two population sizes *N* = 50 and 200, with *L* = 5 parents. The sequencing design aimed to study the effect of read depth *D* and the number *M* of markers for three diallel designs with *L* = 2, 5, and 10 parents, the number of offspring per parent being fixed to 90 so that *N* = 180, 450, and 900, respectively.

To study the robustness to errors in the input marker map, we first simulated SNP array data in the diallel design with no selfings (*S* = 0) and *L* = 5 parents for two population sizes *N* = 50 and 200, using the true parental haplotypes with *M* = 242 markers. To study the effect of markers that are wrongly positioned in long range, we disturbed marker ordering by randomly selecting *f_exch_M*/2 markers on one chromosome arm and *f_exch_M*/2 markers on the other arm, and then exchanging them between two arms. To study the effect of erroneous local marker ordering, we obtained a disturbed genetic map by ordering markers according to the sum of true marker index *t* and a normal distributed random variable with mean 0 and standard deviation *σ_local_*, while keeping the original marker locations.

#### Real Potato datasets

A set of 32 chromosome-length SNP haplotypes from potato were used as the true parental haplotypes to simulate populations and evaluate algorithm performance; see Supplementary Material, Table S1. The 32 haplotypes correspond to chromosome group 4 of 8 tetraploid potato clones, genotyped with version 2 of the potato SNP array, which had 12K markers (Hamilton *et al.* 2011; Felcher *et al.* 2012). The eight clones were mated in pairs to create four F1 populations (Endelman *et al.* 2018), and the software MAPpoly (Mollinari and Garcia 2019; Mollinari *et al.* 2020) was used for parental phasing.

In addition, a 3×3 half-diallel population in potato was used for evaluation; see Table S2 for the dosage data with physical map, and Table S3 for the mating design. Three parents (W6511-1R, W9914-1R, and Villetta Rose) were mated in all three pairwise combinations to create a total population of 434 clones (individual F1 population sizes of 162, 155, and 117). Clones were genotyped with version 3 of the potato SNP array, which had an additional 9K markers from Vos *et al.* (2015) compared to version 2. Allele dosage was assigned using R package fitPoly (Voorrips *et al.* 2011; Zych *et al.* 2019) and 5078 markers distributed across all 12 chromosome groups remained after curation. Physical positions for the input map were based on the potato DMv4.03 reference genome (Potato Genome Sequencing Consortium 2011; Sharma *et al.* 2013).

#### Parameter setup

For simulated data, local ordering and inter-marker distances were refined only when studying the robustness to errors in the input genetic map. For real potato data, PolyOrigin estimated the inter-marker distances, conditional on the input marker ordering. We set up TetraOrigin to have the same option values as those of PolyOrigin. We set up MAPpoly by following its online tutorial. See the Supplementary Materials for the detailed description of the parameter setup for running PolyOrigin, TetraOrigin, and MAPpoly.

### Data availability

PolyOrigin has been implemented in Julia 1.5.3, and is freely available under the GNU General Public License 3.0 from the web site: https://github.com/chaozhi/PolyOrigin.jl. Real potato datasets in Tables S1-S3 are available at FigShare.

## Results

### Comparisons with previous methods

Figure 2 shows the comparisons of PolyOrigin with TetraOrigin and MAPpoly for a single F1 population considering quadrivalent formation (double-reduction is possible). As shown in Figure 2A, both PolyOrigin and MAPpoly have no phasing error when population size *N* ≥ 100, but the MAPpoly software did not produce a solution for the smaller sizes, whereas the phasing error for TetraOrigin was around 0.02 because of the parental dosage errors in the simulated data (ε = 0.01). PolyOrigin and MAPpoly deleted those markers with parental dosage errors (Figure 2B), while TetraOrigin has no function of marker deletion. Note that TetraOrigin may account for parental dosage errors by assuming a non-zero parental genotyping error probability, but this leads to much longer computation time.

**Figure 2:**
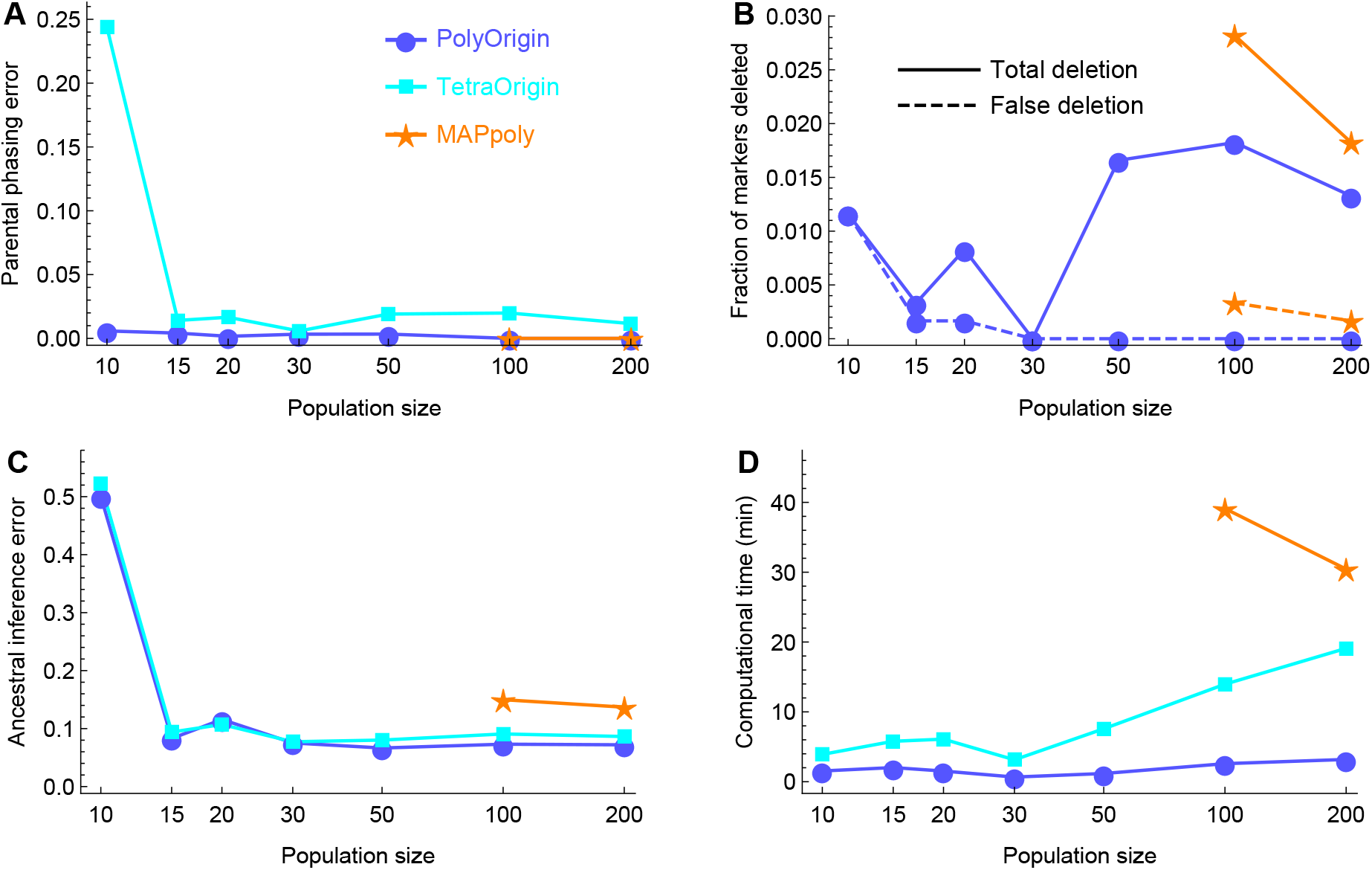
Comparisons of PolyOrigin, TetraOrigin, and MAPpoly in a single F1 population considering quadrivalent formation (double reduction is possible). (**A&C**) Errors in parental phasing and ancestral inference, respectively. (**B**) Fraction of markers deleted. The input number of markers *M* = 201. TetraOrigin has no marker deletion. The dashed lines denote the fraction of markers that are deleted and have no parental dosage errors. (**D**) Computational time in minutes.

Figure 2C shows that TetraOrigin has slightly worse performance in ancestral inference than PolyOrigin, resulting from its higher parental phasing error (Figure 2A). On the other hand, the worse performance of MAPpoly than TetraOrigin and PolyOrigin is mainly because MAPpoly does not account for double reduction. Figure S1 shows that MAPpoly has a similar parental phasing error and a lower ancestral inference error for the simulated data without double reduction.

Figure 2D shows that the computational time of TetraOrigin is around 6 times as long as that of PolyOrigin for population size *N* = 200, although the algorithm of PolyOrigin is almost the same as TetraOrigin for a single F1 population. In comparison, MAPpoly is around 10 times as long as that of PolyOrigin for *N* = 200. For the smaller population sizes (*N* ≤ 50), MAPpoly collapsed for unknown reasons.

### Effect of population design

Figure 3 shows the effect of population design on parental phasing, where the effect of the four design parameters: mating design, population size *N*, number *S* of selfings, and number *L* of parents, is summarized through the number of gametes contributed by each parent. Note that the number of gametes is the same as the number of offspring produced by each parent in the case of no selfings (*S* = 0). It is shown that the parental phasing error becomes very small (<0.01) when the number of gametes from each parent is no less than 30. One exception out of 792 data points in Figure 3A is the high phasing error 0.1 at the number 50 of gametes, corresponding to the middle parent in one of three replicate datasets with linear design, *L* = 3, *S* = 0, and *N* = 50. Further examination shows that the exceptional high error results from a single switch error in the parental haplotypes.

**Figure 3:**
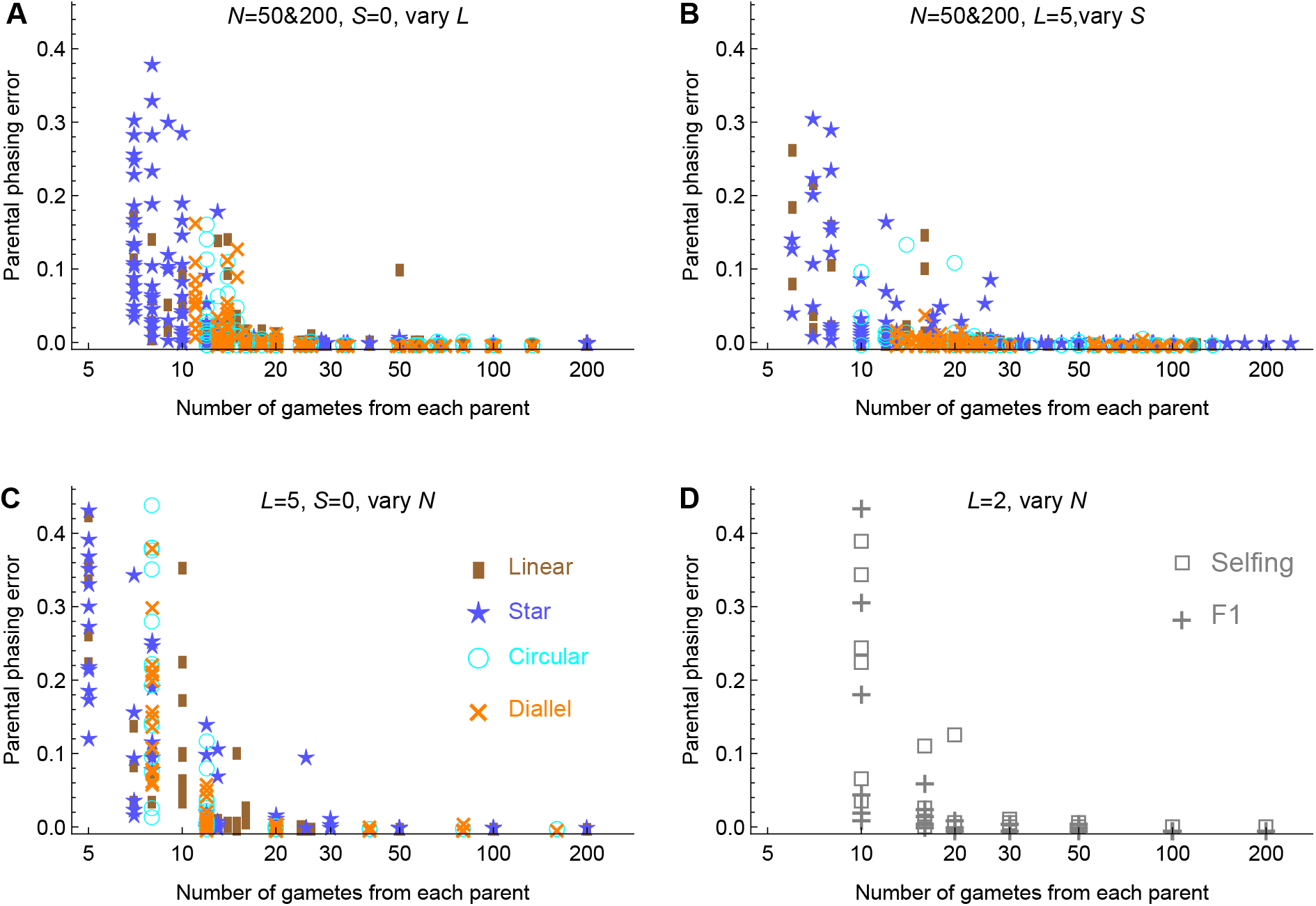
Effect of population design on parental phasing. The x-axis denotes the number of gametes contributed by each parent. The y-axis denotes the parental phasing error for each parent in each of the three replicates given each combination of the design parameter values. (**A-C**) Effect for the populations with varying number *L* of parents, number S of selfings, and population size N, respectively, for each of the four mating designs. Panels **A-B** include the results for two sizes of 50 and 100. (**D**) Effect for bi-parental F1 and two independent selfing populations.

Figure S2 shows the effect of the four design parameters on parental phasing, where the phasing error is averaged over parents and replicates for a given combination of the four design parameter values. It is not unexpected that the parental phasing error increases with the number *L* of parents and decreases with the total population size *N*. For the small population size *N* ≤ 50, the star mating design performed much worse than the linear, circular and diallel designs, particularly at the medium number *S* of selfings, where the numbers of gametes contributed by parents are more unequal that at the two extreme values of *S*. Figure S2F shows that there are no noticeable differences between a single F1 population of size N and the collection of two independent selfing populations of size *N*/2; see also Figure 3D.

Figure S3 shows that the effect of population design on ancestral inference mainly results from its effect on parental phasing.

### Effect of genotyping design

#### SNP array design

Figure 4A, C, and E show the effect of dosage error probability *ε* in the diallel populations with population sizes *N* = 50 and 200. Figure 4A and C show that PolyOrigin is robust to *ε*, except for small *N* = 50 and large *ε* > 0.1, and Figure 4E shows that the fraction of markers deleted increases gradually with *ε* but it is always smaller than *ε*, indicating that both marker deletion and parental error correction contribute to the robustness.

**Figure 4:**
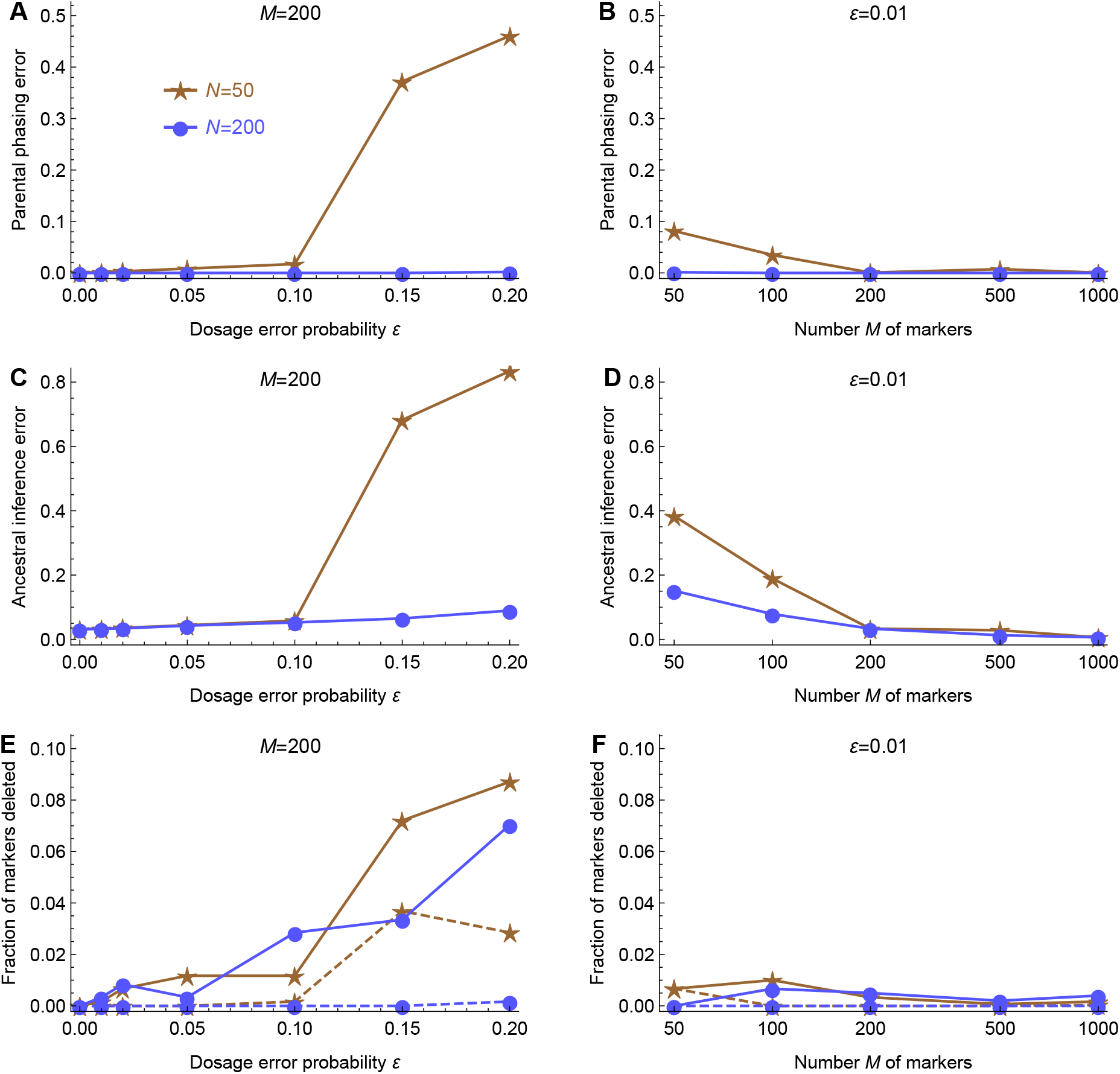
Effect of dosage error probability *ε* and marker density for SNP array dosage data in the diallel populations with no selfing (*S* = 0) and *L* = 5 parents. (**A**, **C** & **E**) Effect of *ε* on parental phasing, ancestral inference, and marker deletion, respectively, with *M* = 200. (**B**, **D** & **F**) Effect of marker density on parental phasing, ancestral inference, and marker deletion, respectively, with *ε* = 0.01. The dashed lines in (**E** & **F**) denote the fraction of markers that are deleted and have no parental dosage errors.

Figure 4B, D, and F show the effect of marker density. Figure 4B shows that parental phasing is robust to marker density except for small *N* = 50 and low *M* ≤ 100, and Figure 4D shows that the ancestral inference error decreases rapidly with marker density. Figure 4F shows that the fraction of markers deleted is independent of marker density and is always smaller than ε.

#### Sequencing design

Figure 5 shows the effect of read depth *D* (nunber of reads per marker per individual) and the number *M* of markers for NGS data in the diallel populations with *L* = 2, 5, and 10 parents, the total population size *N* being adjusted so that the number *N/L* of offspring per parent is fixed. Figure 5A and C show that parental phasing is robust to read depth and marker density, except for low *D* < 10 and small *M* < 250. As shown in Figure 5B and *D*, the ancestral inference error decreases with *M* up to 2000 and with *D* up to 20, and it levels off when *D* > 20.

**Figure 5:**
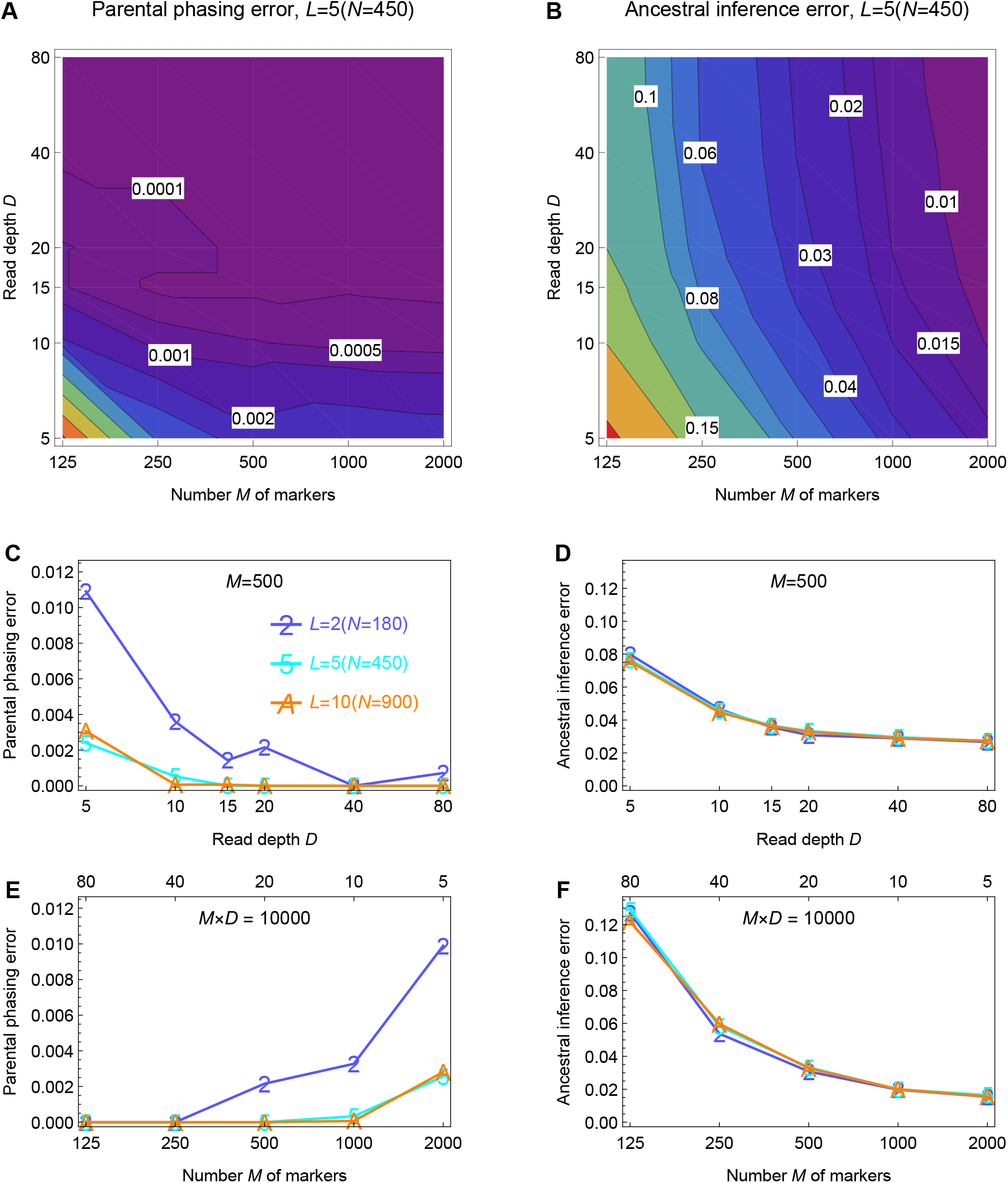
Effect of read depth *D* and the number *M* of markers for NGS data in the diallel populations with no selfing (*S* = 0) and *L* = 2, 5, and 10 parents. (**A**) Contour plot of the parental phasing error as a function of the number *M* of markers and read depth *D*. (**B**) Contour plot of the ancestral inference error as a function of *M* and *D*. (**C&D**) Effect of read depth *D* on parental phasing and ancestral inference, respectively, with *M* = 500. (**E&F**) Effect of read depth *D* on parental phasing and ancestral inference, respectively, with *M* × *D* = 10000.

Figure 5E and F show the effect of *D* and *M*, under the constraint that *D* × *M* = 10000, where the product *D* × *M* denotes the total number of reads, or the NGS cost per individual. Figure 5F shows that the optimal strategy for decreasing ancestral inference error is to increase *M* instead of *D* under the cost constraint, although parental phasing error increases with *M* but it is still very small at *M* = 2000 or *D* = 5 (Figure 5E).

Figure 5C-F show that the number *L* of parents has little effect on parental phasing and ancestral inference, if the population size *N* is increased proportionally, although the parental phasing error for *L* = 2 is slightly greater than that for *L* = 5 and 10.

### Robustness to errors in input map

Figure 6 shows map refinement in the presence of long-range or local disturbances in the input genetic maps in the diallel populations with population sizes *N* = 50 and 200. Figure 6A-B show that map improvement is more effective in the large populations (*N* = 200) than in the small populations (*N* = 50), and that it is more effective in the presence of long-range disturbances that in the presence of local disturbances. This is because most markers with long-range disturbances have been deleted (Figure 7E), while few markers with local disturbances have been deleted (Figure 7F). Figure 6C-D show that map length is slightly underestimated and inflated under strong disturbances.

**Figure 6:**
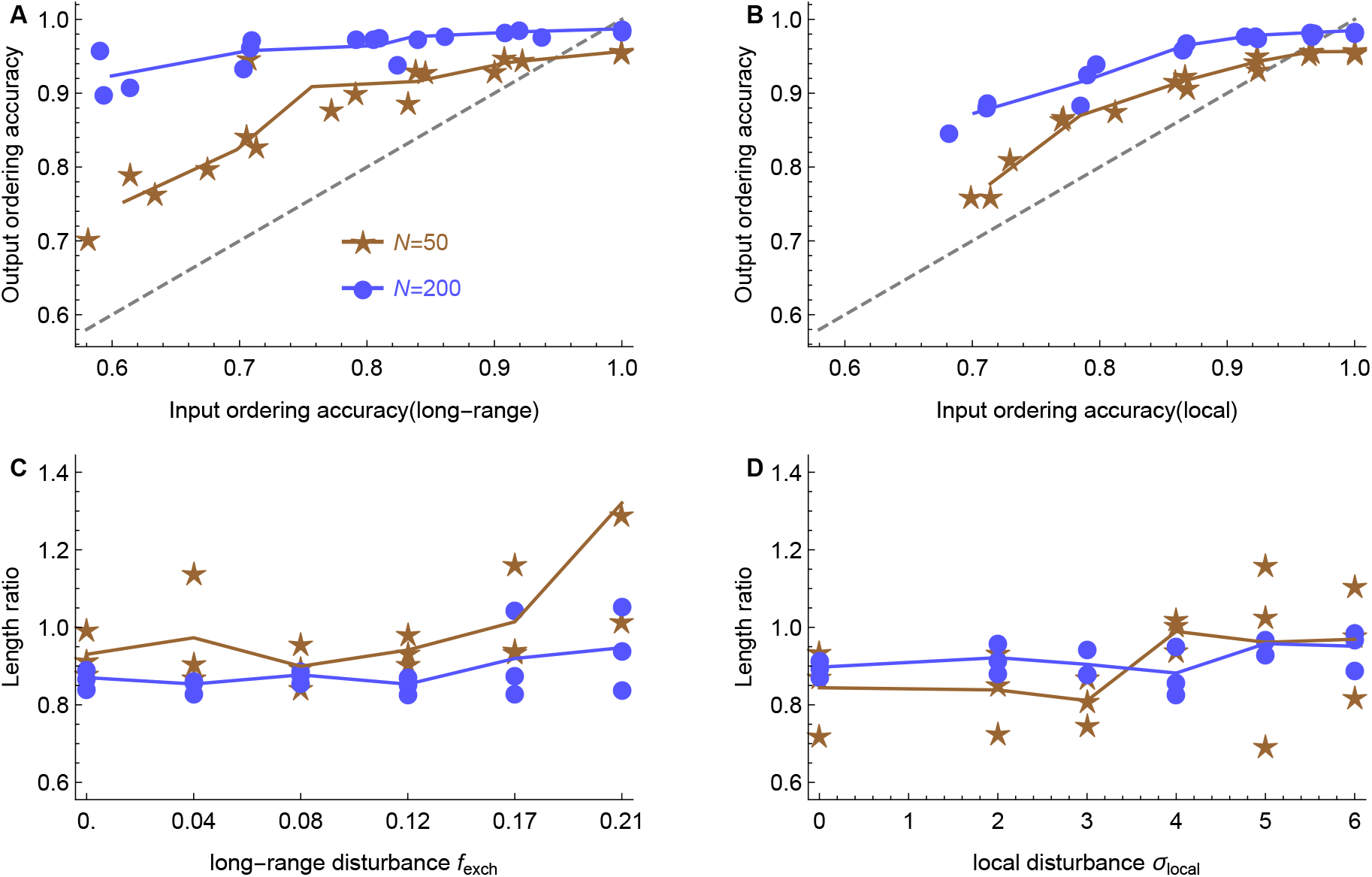
Refinement of the input genetic maps with long-range or local disturbances in the diallel populations with no selfings (*S* = 0) and *L* = 5 parents. (**A&B**) Improvement of marker ordering in the presence of long-range and local disturbances, respectively, the dashed lines denoting *y* = *x*. (**C&D**) Ratio of estimated genetic length to true value in the presence of long-range and local disturbances, respectively.

**Figure 7:**
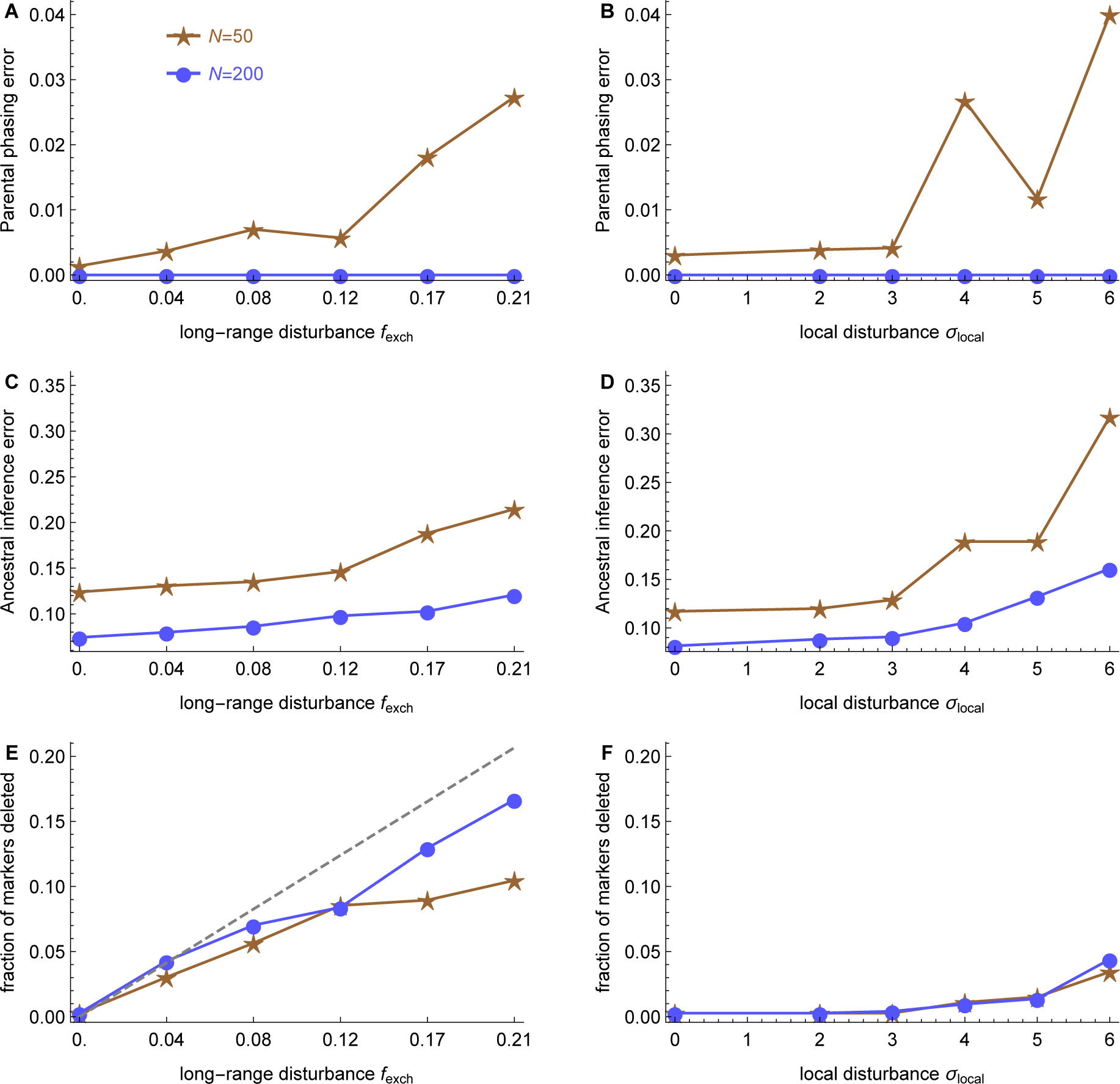
Effect of long-range or local disturbances in the input genetic maps in the diallel populations with no selfings (*S* = 0) and *L* = 5 parents. The left and right panels denote the effect of long-range and local disturbances, respectively. (**A&B**) Effect on parental phasing. (**C&D**) Effect on ancestral inference. (**E&F**) Fraction of markers deleted. The dashed line denotes *y* = *x*.

Figure 7 shows that both parental phasing and ancestral inference are robust to long-range or local disturbances in the input marker maps, although the ancestral inference error slightly increases with the disturbance strength. Figure 7A-D show that the robustness is stronger in large populations (*N* = 200), partially because marker deletion and parental error correction are less effective in small populations (*N* = 50).

### Evaluation with real data

PolyOrigin was applied to a 3 × 3 half-diallel population of autotetraploid potato. The inferred frequency of quadrivalents is 19% on average, ranging from 9% to 40% across the 12 chromosomes, and the frequencies of the three possible bivalent pairings for each parent were nearly equal, as expected for a true autopolyploid (Figure S4). Of the 5078 markers, 32 were discarded due to poor fit, and 11 genotype errors were detected in the parents, 10 of which involved an allele dosage error of magnitude 1. Even though all 434 progeny had passed sensitive quality control tests for parentage based on the genome-wide markers (ENDELMAN *et al.* 2017), PolyOrigin flagged 19 outlier offspring due to an excessive number of haplotype breakpoints (Figure S5).

Double reduction refers to the inheritance of both sister chromatids at a single locus in the diploid gamete. Figure 8A shows one such offspring, and the double reduction events are visible as dark blue segments in linkage groups 2, 5, and 6. The predicted haplotypes from MAPpoly (Figure 8B) are similar to PolyOrigin except in regions of double reduction, where the MAPpoly solution tends to shows a large number of haplotype breakpoints (Figure S5). Figure 8C shows that the fraction of gametes with double reduction obtained by PolyOrigin increases from almost 0 at centromeres to the maximum 0.078 at telomeres. Note that the fraction would increase by a factor of about 2 if it had been calculated as the faction of zygotes with double reduction (Bourke *et al.* 2015).

**Figure 8:**
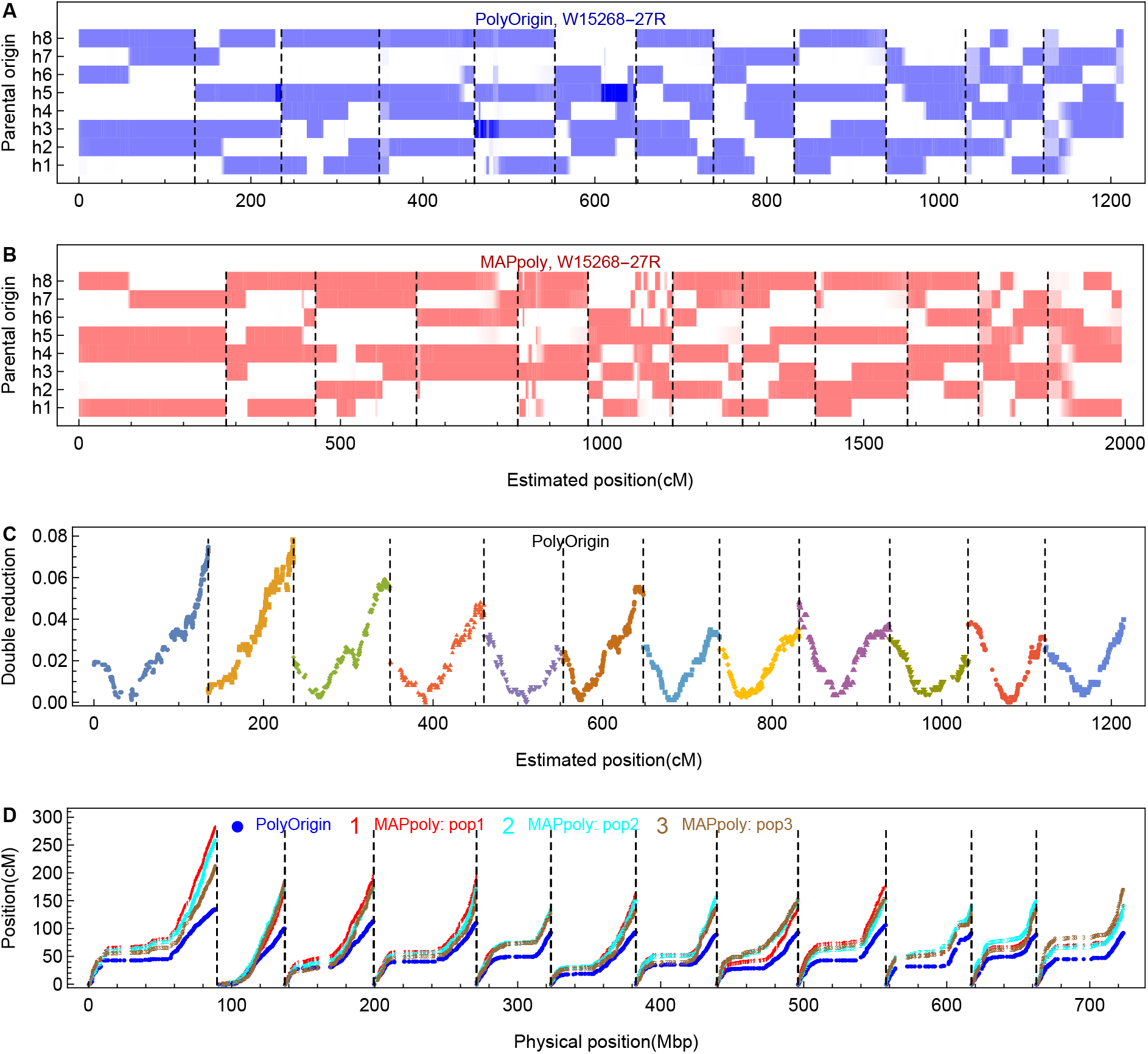
Comparison of PolyOrigin with MAPpoly for the 3×3 potato diallel population. Dashed vertical lines denote chromosome boundaries. (**A**) Posterior probabilities obtained by PolyOrigin for the example offspring (W15268-27R). The darker the color, the higher the probability. (**B**) Posterior probabilities obtained by MAPpoly for the same example offspring. (**C**) Variation of double reduction along chromosome obtained by PolyOrigin. The y-axis denotes the fraction of gametes having two copies of the same parental haplotype, based on the maximum possible origin-genotypes of offspring at a given marker. (**D**) Comparison of the estimated genetic maps with the physical map. On the y-axis of (**A&B**), h1-h4 denote the homologs from the first parent (W6511-1R) of the offspring, and h5-h8 for the second parent (VillettaRose).

Another notable difference between the PolyOrigin and MAPpoly solutions is the length of the genetic map (Figure 8D). The MAPpoly map was 19.4 Morgans (M) compared to 12.1 M for PolyOrigin, which is more similar to the estimates of 10–11 M published in biparental linkage mapping studies (Massa *et al.* 2015; Bourke *et al.* 2016; Da Silva *et al.* 2017). One source of map inflation with MAPpoly appears to be elevated estimates of recombination frequency in the pericentromeric regions (Figure 8B). Even when the three F1 populations were analyzed separately with PolyOrigin, more accurate map lengths were obtained (Figure S6)

Similar to the simulation studies, PolyOrigin was much faster than MAPpoly in analyzing the real potato data (*N* = 434 and *M* = 5078). The computational times were 230 hours for MAPpoly, 10 hours for PolyOrigin analyzing the three F1 populations jointly, and 4.9 hours for PolyOrigin analyzing the data separately. We did not use parallel computation in the analysis, although both PolyOrigin and MAPpoly can perform parallel computation at the chromosome level.

## Discussion

We have developed a new method, implemented in PolyOrigin, for haplotype reconstruction in connected tetraploid F1 populations, each F1 population being produced by cross-fertilization between two parents or self-fertilization from a single parent. PolyOrigin extends the previous HMM framework TetraOrigin (Zheng *et al.* 2016) from a F1 cross to multiple F1 crosses. Both PolyOrigin and TetraOrigin use a forward-backward procedure for parental phasing, whereas MAPpoly (Mollinari and Garcia 2019; Mollinari *et al.* 2020) uses only a forward procedure for parental phasing in a F1 cross. This algorithmic difference may explain why MAPpoly did not work for small population sizes.

In comparison to the basic steps of parental phasing and ancestral inference in TetraOrigin, PolyOrigin has added a procedure of marker deletion in the step of parental phasing. The marker deletion is based on the Vuong’s closeness test (Vuong 1989) with the default significant level 0.05, which has been shown to be very effective to remove long-range misplaced markers and some markers with parental errors. In the parental phasing by sequentially adding markers, MAPpoly uses two limit parameters controlling marker deletion: one for the maximum increase of map length, and one for the maximum number of linkage phase configurations to be tested. It is not obvious how to set these parameter values, and too many testing phase configurations will considerably increase computation time.

PolyOrigin has also added a procedure of parental error correction in the step of ancestral inference. The procedure corrects parental dosages and phases by minimizing the number of mismatches between the observed and estimated genotypes in offspring, conditional on phased parent genotypes, which is computationally more efficient than TetraOrigin introducing a parental dosage error parameter. Not surprisingly, the error correction procedure is not effective in small populations, particularly, with low depth NGS data.

Another quality-control feature implemented in PolyOrigin is the automated outlier detection of progeny with an excessive number of haplotype switches. In the simulated datasets, very few outliers were ever detected, which suggests a very small false discovery rate. However, we are unable to explain why 19 of the 434 potato progeny were outliers. The potato SNP array has been shown to be a powerful tool for detecting pedigree errors (Endelman *et al.* 2017), and all 434 progeny passed these quality control measures. Perhaps some of the complex chromosomal behavior possible in meiosis I is poorly captured by the genetic model in PolyOrigin. The average frequency of 27% quadrivalents in the potato population, with some variation between parents and chromosomes, is consistent with previous studies based on marker data (Bourke *et al.* 2015) and cytological techniques (Choudhary *et al.* 2020).

To increase the robustness to dosage uncertainties in low depth NGS data, PolyOrigin has integrated a dosage calling procedure by analyzing read counts directly, where the probabilities of read counts givens all possible dosages are calculated. These probabilities can also be provided by posterior dosage probabilities exported by the softwares such as polyRAD (Clark *et al.* 2019) for NGS data and fitPoly (Voorrips *et al.* 2011; Zych *et al.* 2019) for SNP array data. in comparison, TetraOrigin can analyze only dosage data, and MAPpoly cannot analyze read counts directly, relying instead on an input file with genotype probabilities.

PolyOrigin allows flexibility in the mating and genotyping designs for linkage mapping projects. Our results show that the parental phasing error is less than 0.01 when the number of offspring per parent is over 30. This implies that incomplete diallel designs, such as linear or star, can be used with similar performance to a complete diallel, which can be difficult to create due to reproductive limitations of the parents. We also show that because PolyOrigin effectively pools data across the entire chromosome, reliable genotype calls can be made in autotetraploids with much less read depth per marker, such as 10 or 20X, compared with values of 60-80X when genotype calls are made independently for each marker (Uitdewilligen *et al.* 2013; Matias *et al.* 2019). For the design of sequence-based genotyping platforms with a fixed number of markers (e.g., baits or amplicons) and reads per sample, we have shown that increasing the number of markers leads to more accurate results even though the number of reads per marker decreases.

Computationally, PolyOrigin is about one order of magnitude faster than TetraOrigin, mainly because TetraOrigin is implemented in Mathematica (Wolfram Research 2016) while PolyOrigin is implemented in Julia (Bezanson *et al.* 2017). Although MAPpoly is implemented in R (R Core Team 2019) and C/C++, it is more than one order of magnitude slower than PolyOrigin, probably because the phasing algorithm of MAPpoly requires two-point linkage analyses. in addition, the computational time of PolyOrigin scales linearly with the number of parents, population size, and the number of markers (Figure S7).

PolyOrigin has been implemented and tested for tetraploid, and most parts of the algorithm can be extended easily to higher ploidy levels. However, a stochastic algorithm would be needed to infer valent formations for hexaploids or higher, because the number of possible valent formations increases rapidly with ploidy level and the current implementation of PolyOrigin considers all possible configurations. For example, there are 105 possible bivalent chromosome pairings in octoploid and thus 105^2^ combinations for biparental populations, not to mention the demanding modeling and computational requirements for multivalent formation.

In conclusion, we have developed a novel method PolyOrigin for haplotype reconstruction in connected tetraploid F1 populations, which opens up exciting new possibilities for haplotype-based QTL mapping in such populations. Extensive evaluations have shown that PolyOrigin is robust to various sources of errors in input genetic data and is around one order of magnitude faster than the previous methods that works only for a single F1 population.

## Acknowledgments

Financial support provided by USDA NIFA Award No. 2019-67013-29166. CZ developed the PolyOrigin model and algorithm and wrote the first draft of the manuscript. RRA wrote the wrap-up PedigreeSimR and helped using MAPpoly. JBE provided potato data and helped reshape the manuscript. All authors participated in study design and provided critical feedback. All authors read and approved the final manuscript.

## Supplementary Materials

### Parameter setups

#### PolyOrigin

For a simulated dataset, the Julia command line used for PolyOrigin is given by

~~~
polyOrigin(genofile, pedfile)
~~~

where genofile specifies input marker data, including genetic map, and genotypic data of parents and offspring, and `pedfile` specifies the population mating design. The default settings *epsilon=0.01* and *seqerr = 0.001* are used, specifying the initial value for the interal estimation of dosage error proability and the sequencing error probability in the case of read count data. By default, the input marker map is genetic map and it will not be refined (*isphysmap=false),* parental phasing assumes only bivalent formations (*chrpairing_phase=22),* and both bivalent and quadrivalent formations are considered for ancestral inference and parental error correction (*chrpairing=44).*

For the real potato dataset with physical map, the Julia command line is given by

~~~
polyOrigin(genofile,pedfile,
    isphysmap=true, recomrate=1.25,
    refinemap=true, refineorder=false)
~~~

where the keyword argument *isphysmap* specifies that input map is physical map with marker positions in unit of base pair, and recomrate specified the global constant recombination rate in unit of cM/Mbp. *refinemap=true* indicates the performance of map refinement, and *refineorder=false* indicates the refinement of inter-marker distances but not marker ordering.

### TetraOrigin

The Mathematica command line used for TetraOrigin is given by

~~~
inferTetraOrigin[genofile, epsO, epsF, ploidy, outstem,
    maxStuck -> 5, maxIteration -> 30, maxPhasingRun -> 10,
    bivalentPhasing -> True, bivalentDecoding -> False]
~~~

where `genofile` specifies the input genotypic data. *epsF* and *epsO* specify the dosage error probability in parents and offspring, respectively. *ploidy=4* for tetraploids, and *outsem* specifies the string ID of output file. The options maxStuck, maxIteration, and maxPhasingRun for the parental phasing algorithm are re-set to be consistent with PolyOrigin. And the default settings for bivalentPhasing and bivalentDecoding are consistent with PolyOrigin.

For the simulated F1 datasets, we set *epsO* to the true value 0.01. Although the true parental error probability is also 0.01, we set *epsF=0* because a non-zero setting would result in much longer computational time.

### MAPpoly

We closely follow the online MAPpoly tutorial on building a genetic map using potato genotype data. The R command lines used for MAPpoly are divided into the following steps

~~~
#step1: read data
dat.dose.csv <-read_geno_csv(file.in = genofile, ploidy = 4)

#step2: marker filtering
pval.bonf <-0.05/dat.dose.csv$n.mrk
dat.chi.filt <-filter_segregation(dat.dose.csv,
    chisq.pval.thres = pval.bonf, inter = FALSE)
dat.seq <-make_seq_mappoly(dat.chi.filt, “all”)

#step3: two-point analysis
counts <-cache_counts_twopt(input.seq = dat.seq, get.from.web = TRUE) all.rf.pairwise <-est_pairwise_rf(input.seq = dat.seq,
    count.cache = counts, n.clusters = 1)

#step4: parental phasing and marker spacing for a given marker ordering map <-est_rf_hmm_sequential(input.seq = dat.seq,
   start.set = 10,
   thres.twopt = 10,
   thres.hmm = 10,
   extend.tail = NULL,
   info.tail = TRUE,
   twopt = all.rf.pairwise,
   sub.map.size.diff.limit = 20,
   phase.number.limit = 50,
   reestimate.single.ph.configuration = TRUE,
   tol = 10e-3,
   tol.final = 10e-4)
map.error <-est_full_hmm_with_global_error(input.map = map, error = epsilon))

#step5: calculate genotype probability
genoprob <-calc_genoprob_error(input.map = map.error,
   error = epsilon)
~~~

We skip the step of marker grouping and marker ordering by using the true genetic map or the real physical map. The dosage error probability *epsilon* is set to the true value for simulating data, and 0.02 for the real potato data, based on the estimation of PolyOrigin.

## Supplementary figures

**Figure S1:**
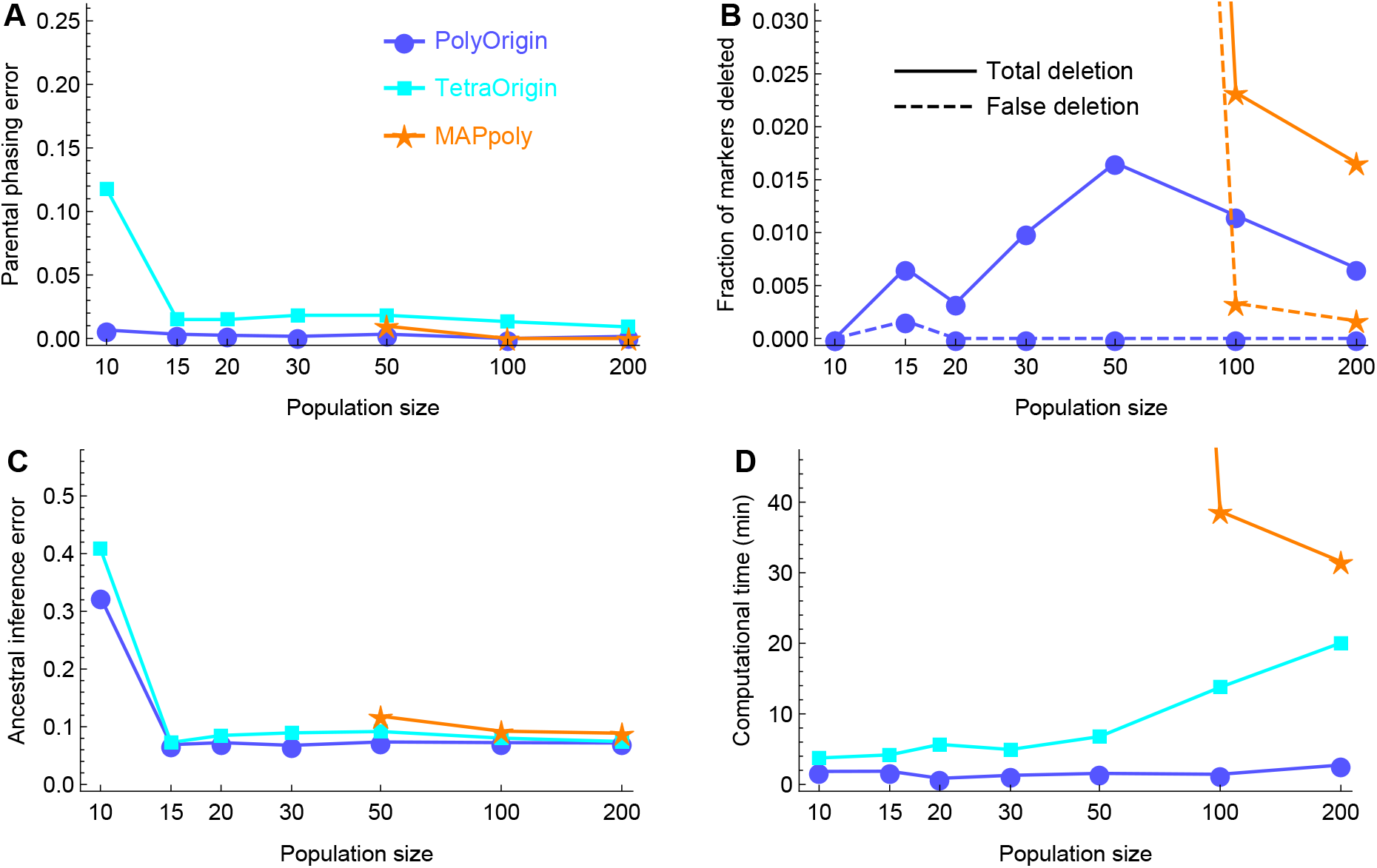
Comparison of PolyOrigin, TetraOrigin, and MAPpoly for the simulated F1 populations without double reduction. The dashed lines in (**C**) denote the fraction of markers that are deleted and have no parental dosage errors. For *N* = 50, MAPpoly deleted 23% markers and took the computational time of 303 minutes.

**Figure S2:**
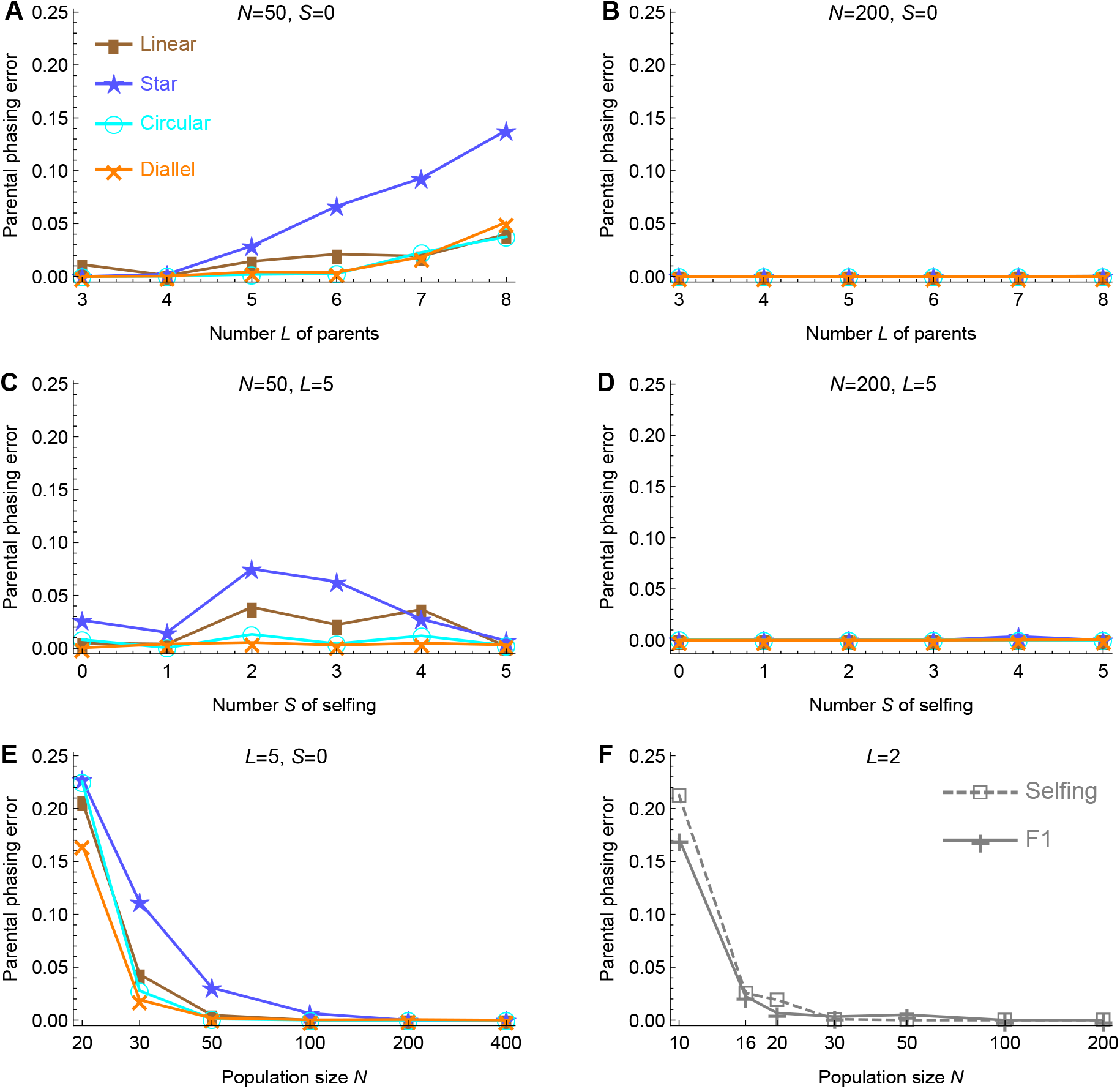
Effect of population design on parental phasing. (**A&B**) Effect of the number *L* of parents for populations with no selfings (*S* = 0) and sizes of *N* = 50 and 200, respectively. (**C&D**) Effect of the number *S* of selfings for populations with *L* = 5 parents and sizes of *N* = 50 and 200, respectively. (**E**) Effect of population size *N* for *L* = 5 parents. (**F**) Effect of population size *N* for bi-parental F1 and two independent selfing populations.

**Figure S3:**
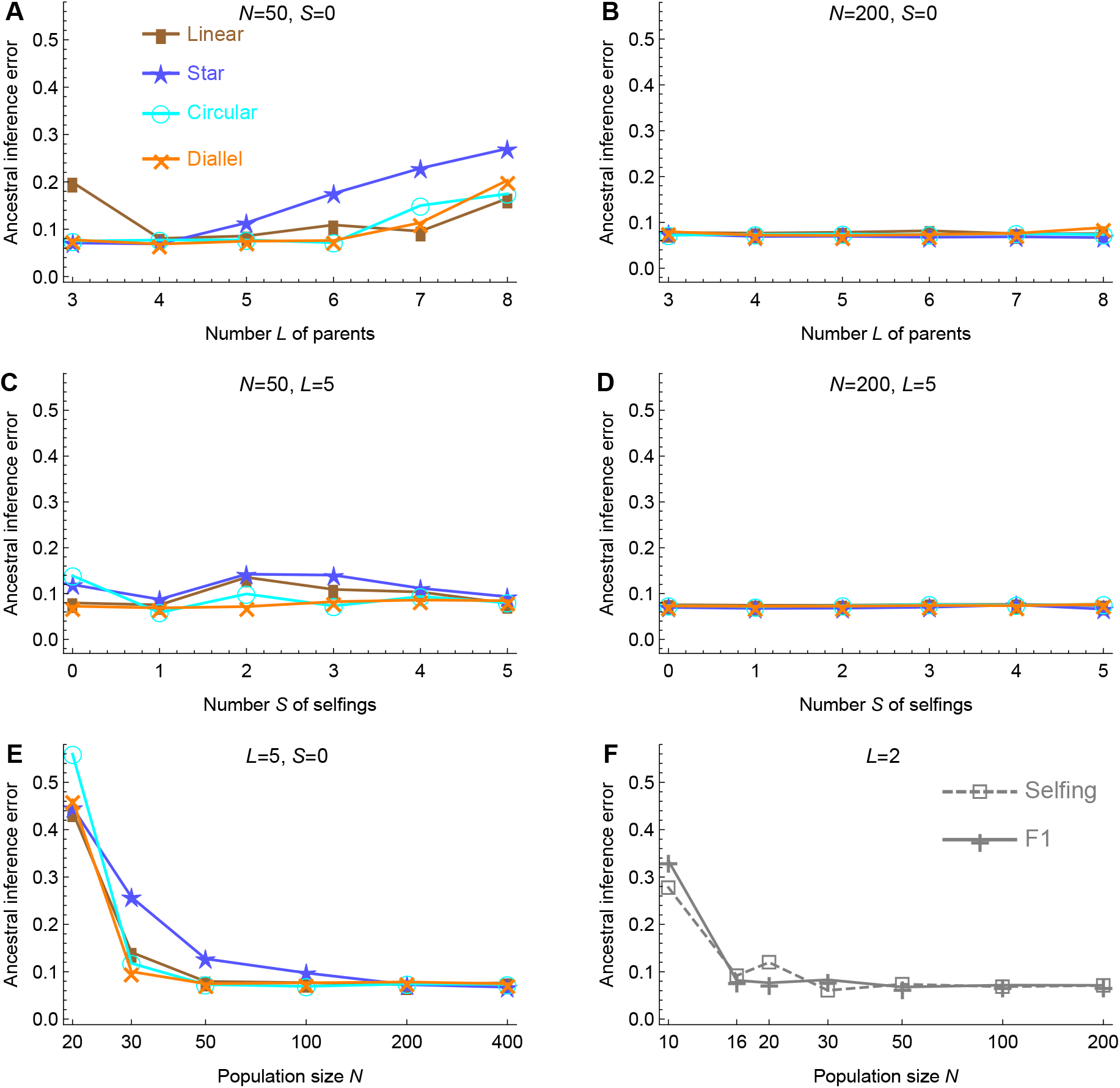
Effect of population design on ancestral inference. (**A&B**) Effect of the number *L* of parents for populations with no selfings (*S* = 0) and sizes of *N* = 50 and 200, respectively. (**C&D**) Effect of the number *S* of selfings for populations with *L* = 5 parents and sizes of *N* = 50 and 200, respectively. (**E**) Effect of population size *N* for *L* = 5 parents. (**F**) Effect of population size *N* for bi-parental F1 and two independent selfing populations.

**Figure S4:**
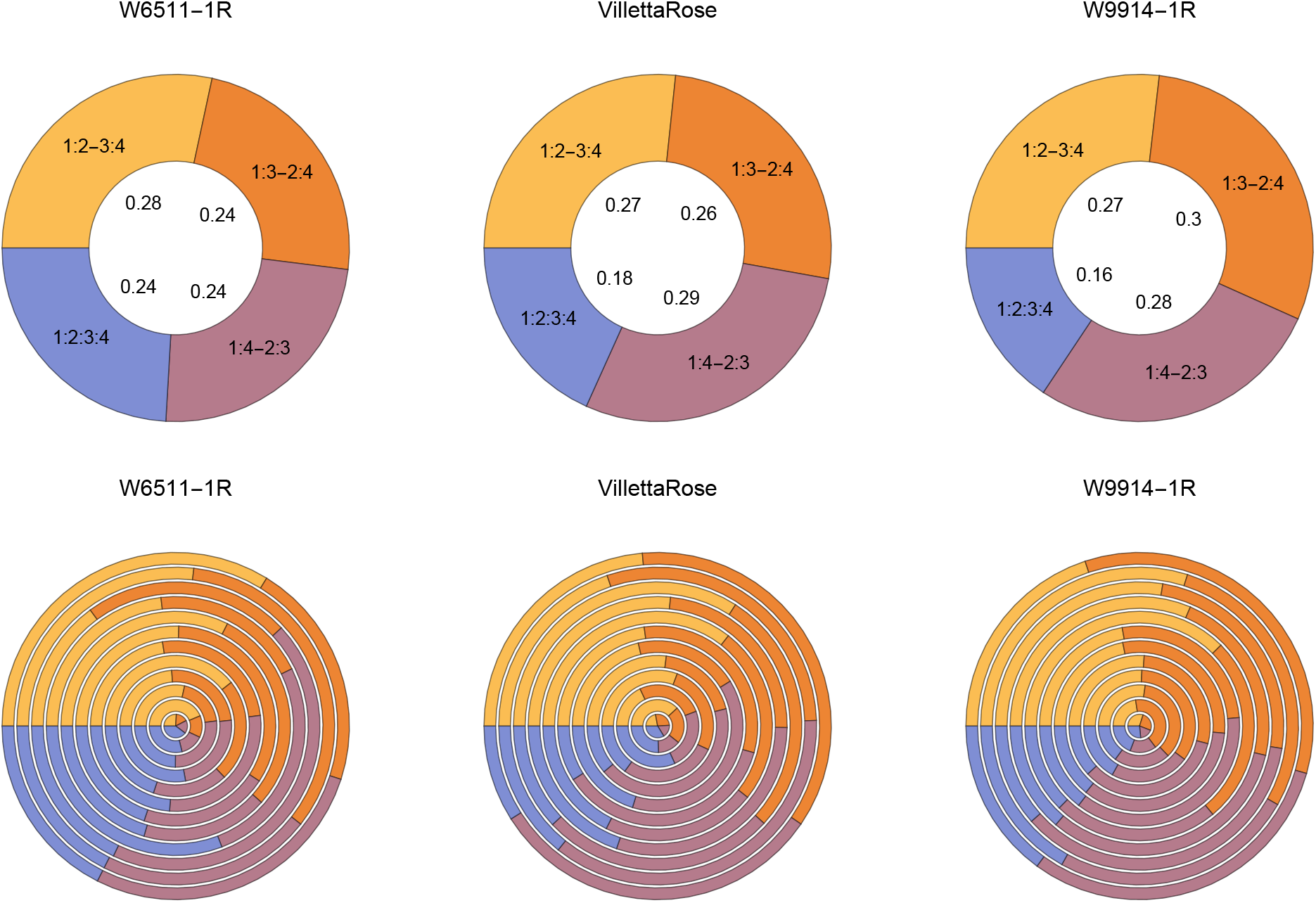
The proportion of valent configurations for the 12 chromosomes of potato in the 3×3 half-diallel with parents VillettaRose, W6511-1R, and W9914-1R. The proportion was calculated based on the maximum possible configurations for each offspring and each chromosome. The configuration 1:2:3:4 refers to a quadrivalent, while the other three refer to bivalent pairs (the colon separates paired homologs). Each bottom panel denotes the proportions among the 12 chromosomes starting from the inner, and the upper panels denote the averages over chromosomes

**Figure S5:**
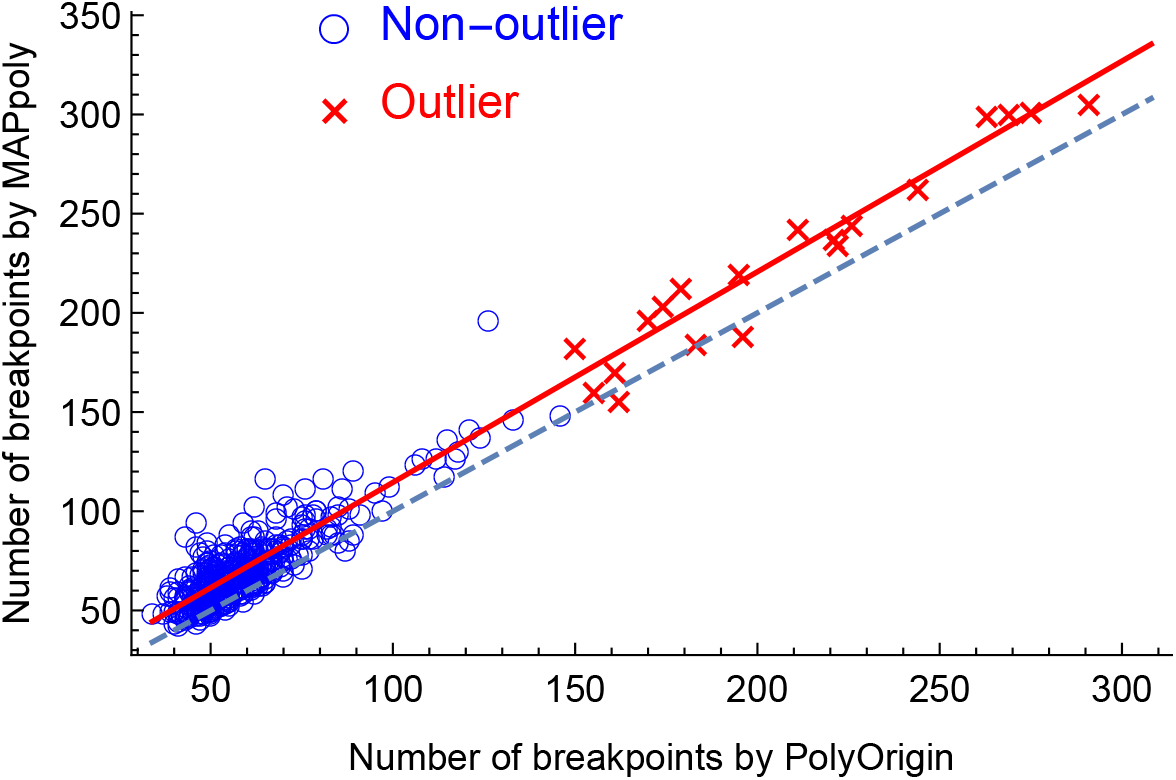
Comparison of PolyOrigin with MAPpoly in terms of the number of haplotype breakpoints for each offspring. Red crosses denote outlier offspring labeled by PolyOrigin, and blue circles denote non-outliers. Dashed line denotes *y* = *x*, and red line denotes the regression line.

**Figure S6:**
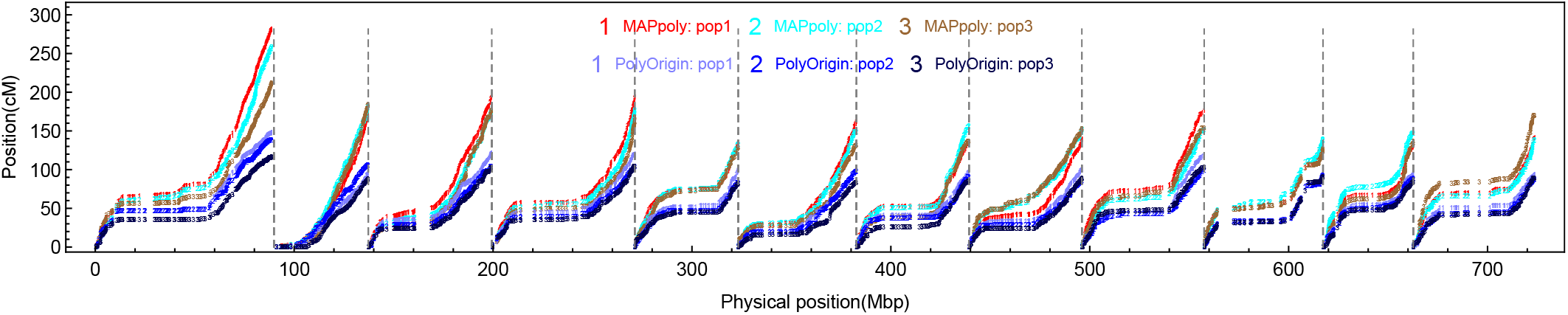
Comparison of PolyOrigin with MAPpoly for each of the three F1 populations in the real 3×3 potato diallel population.

**Figure S7:**
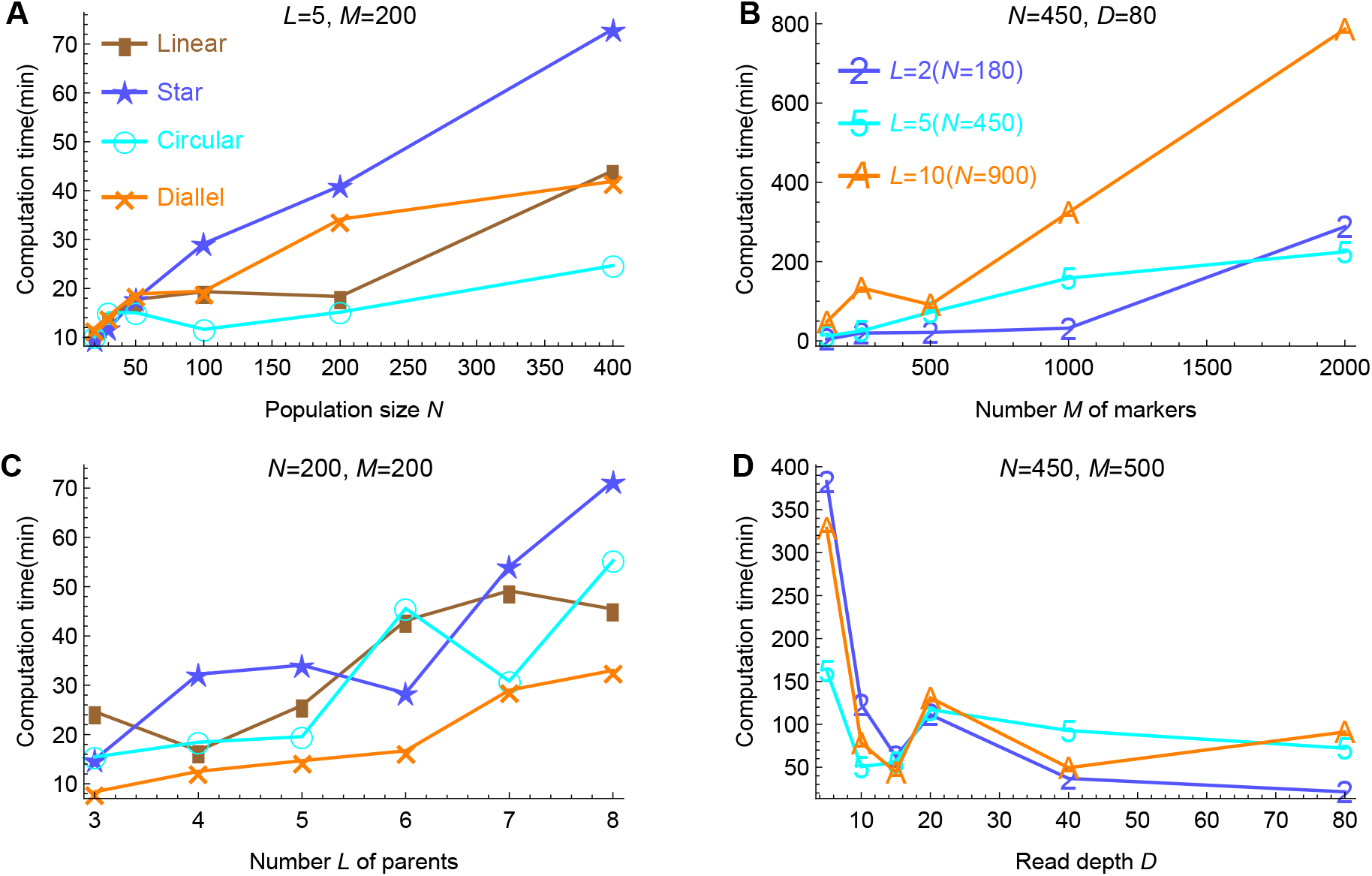
Effect of population design and genotyping design on computational time (in minutes). (**A&C**) Computational time used in analyzing the simulated SNP array data in the four mating designs. (**B&D**) Computational time used in analyzing the simulated GBS data in the diallel design with *L* = 2, 5, and 10 parents, respectively.

